# CFTR function in nasal airway cells from symptomatic and asymptomatic CF heterozygotes

**DOI:** 10.64898/2026.06.17.732907

**Authors:** Audrey Pion, Alyssa Chang, Piero Mejia, Alice C. Eastman, Erin Kavanagh, Ana Topasna, Katherine Starego, Alianna Parr, Karen Raraigh, Christian Merlo, Neeraj Sharma, Garry Cutting

## Abstract

**Rationale:** An estimated 25 million people worldwide have one deleterious variant in the cystic fibrosis transmembrane conductance regulator (*CFTR*) gene. Chronic respiratory disease symptoms are at an increased prevalence in cystic fibrosis (CF) heterozygotes.

**Objectives:** Determine the level of CFTR function in CF heterozygotes compared to individuals without CF-causing variants. Establish whether CFTR function differs between asymptomatic and symptomatic CF heterozygotes.

**Methods:** Individuals without respiratory symptoms or CF family history were recruited as controls. Heterozygotes were recruited from families with a CF individual harboring null alleles or c.1521_1523del (F508del) in *CFTR*. CFTR function was measured by short circuit current in primary human nasal epithelial cells (HNEs) from participants. Cell composition was assessed by single cell RNA sequencing.

**Measurements and Main Results:** CFTR function was variable in cells from control and heterozygous individuals. Mean CFTR function in asymptomatic null (8.8±0.5µA/cm^2^ (SEM); n=30) and F508del (8.7±1.0µA/cm^2^; n=22) heterozygotes was similar and significantly lower at 54.6% and 53.9% than controls (16.1±1.1µA/cm^2^; n=24; p<0.0001). Mean CFTR function in symptomatic heterozygotes (8.4±1.0µA/cm^2^; n =15) was 52.1% of controls and did not differ from asymptomatic heterozygotes (p=0.7803). Cell identities and proportions were equivalent between control and heterozygous cultures. HNEs from CF heterozygotes showed variable response to CFTR modulators.

**Conclusions:** CFTR function in primary airway cells exhibits substantial interindividual variability and overlaps between controls and CF heterozygotes. CF heterozygotes exhibit approximately 50% of CFTR function in controls, regardless of symptom status. These findings suggest that respiratory symptoms in CF heterozygotes are influenced by factors beyond CFTR dysfunction.

**At a Glance Commentary:** *Scientific Knowledge on the Subject:* A growing body of evidence indicates that cystic fibrosis (CF) heterozygotes are at increased risk for a range of common and chronic respiratory diseases. Given that an estimated 10 million individuals in the United States are CF heterozygotes, this population may represent a substantial and underappreciated burden of CFTR-associated disease. However, CFTR function has not been well characterized in CF heterozygotes, and it remains uncertain whether observed clinical phenotypes reflect reduced CFTR activity. Resolving these issues will be essential for clarifying the pathobiology of common respiratory diseases and for evaluating the potential role of CFTR modulator therapy in symptomatic CF heterozygotes.

*What This Study Adds to the Field:* Analysis of 24 controls and 67 CF heterozygotes revealed substantial interindividual variability in CFTR function, as measured *ex vivo* in differentiated nasal airway epithelial cells. Mean CFTR function in CF heterozygotes was approximately 50% of that observed in controls. CFTR function did not differ significantly between symptomatic and asymptomatic heterozygotes. These findings suggest that reduced CFTR activity may contribute to symptom susceptibility in CF heterozygotes, but additional factors beyond CFTR dysfunction are likely required for the development of CF-like features.

*Ethical approval and participant consent statement:* All participants provided written consent to the research study under IRB00116966 and/or IRB00235883 and consented to have their anonymized data published. All research was conducted in a fair and ethical manner.

## Introduction

Cystic fibrosis (CF) is a genetic disorder caused by DNA variants that lead to loss of function of the CF transmembrane conductance regulator (CFTR). The condition is inherited in an autosomal recessive Mendelian pattern, in which a *CFTR* allele bearing a loss of function variant carried by each parent is transmitted to the affected offspring.^1^ Typically, each parent and two-thirds of their unaffected offspring are CF heterozygotes because they carry a dysfunctional and functional *CFTR* allele, the latter of which is thought to protect them from disease development. However, an increased prevalence of chronic conditions that clinically overlap with CF has been reported in CF heterozygotes, thereby challenging the Mendelian concept that heterozygotes do not develop symptoms observed in affected individuals.^2^ These conditions are relatively common in the general population and generally involve the respiratory tract including chronic rhinosinusitis, idiopathic bronchiectasis, allergic bronchopulmonary aspergillosis, and non-mycobacterial tuberculosis. More recently, large-scale retrospective epidemiologic studies documented an increased risk of CF-like complications in CF heterozygotes. An extended list of relevant literature can be found in **Supplemental Table S1**.

These observations raise key questions; what is the level of CFTR function in CF heterozygotes and does reduced CFTR function play a role in individuals manifesting CF-like symptoms? Based on the principles of Mendelian autosomal recessive inheritance and RNA studies, it has been assumed that individuals with one working allele have approximately half the CFTR function as those with two working alleles.^3,4^ Additionally, when two *CFTR* alleles are present *in vivo*, it has been assumed that total CFTR function results from an additive contribution from each allele, but it is possible that synergistic or multiplicative interactions could occur. The answers to the above questions have important healthcare ramifications, as there are estimated to be 10 million CF heterozygotes in the United States^5^ and over 25 million worldwide. Thus, a modest increase in the relative risk for CF-like symptoms in CF heterozygotes could account for a sizeable fraction of individuals with a variety of common chronic respiratory conditions. Furthermore, the expansion of genetic testing in newborn screening, medical clinics, large scale population studies, and direct-to-consumer businesses raises new challenges in the counseling of millions of CF heterozygotes. Finally, success in precision treatment of CF using CFTR targeting drugs might present an opportunity to treat chronic conditions in CF heterozygotes.

To address these questions, we obtained progenitor airway cells from the nasal passages of people without CF-like symptoms (asymptomatic) and no CF-causing variants (“controls”) and from individuals with one CF-causing variant (“CF heterozygotes”) who were either asymptomatic or symptomatic. Cells were proliferated by conditional reprogramming then differentiated into polarized airway cultures to measure CFTR function.^6–9^ This method has been used to establish baseline CFTR function and response to CFTR modulators in nasal cells obtained from individuals with a variety of CF-causing variants.^6,10^ Extensive efforts were made to control for technical variables so the differences in means among groups primarily represented biological differences. This approach allowed us to evaluate the level of CFTR function in asymptomatic CF heterozygotes relative to controls and then compare CFTR function between symptomatic and asymptomatic CF heterozygotes.

## Methods

All materials and methods can be found in the supplemental text.

## Results

### Establishing CFTR function in human nasal epithelial cells (HNEs) from controls

HNEs obtained from the nasal passages of participants were differentiated and expanded into airway cell monolayers (**Figure 1A**) and CFTR function was measured by short circuit current (**Figure 1B**). To evaluate factors that could affect the reproducibility of our culture system, we separately cultured cells isolated from the left and right nostrils of two research participants and determined that CFTR function did not differ at passage 2 (both samples), passage 3 (sample FIN001), or passage 4 (sample FIN001; **Figure S1A**). Similarly, the number of days exposed to PALI media prior to functional testing (**Figure S1B**) did not affect CFTR current measures while the number of 3T3 cells used to support cell growth on Snapwell^TM^ filters had variable effect (**Figure S2)**. In contrast, CFTR currents differed between fresh and cryopreserved cultures (**Figure S3A**), passage number (**Figure S3B**), and number of days cultured at air-liquid interface (**Figure S3C**). As recruitment progressed, we accounted for these variables in as many cultures as possible by performing CFTR functional measurements on fresh samples (i.e. not cryopreserved) at passage 2, differentiated for 24-26 days with a consistent number of 3T3 cells plated beneath Snapwell™ filters during filter expansion and a consistent number of HNEs plated onto Snapwell™ filters.

**Figure 1:**
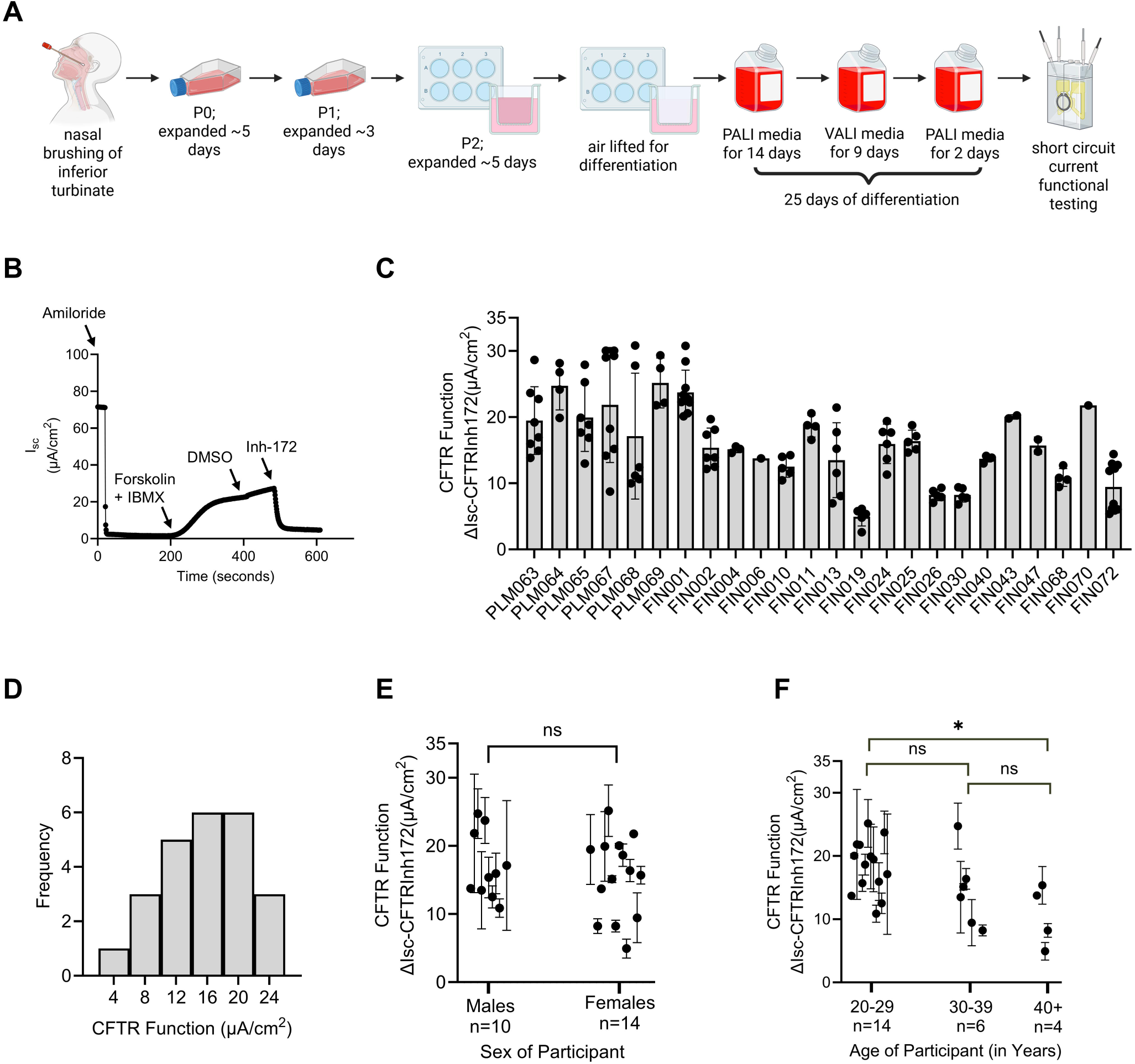
CFTR function in primary nasal airway cells from controls. (A) Diagram illustrating the collection, culturing, and differentiation of human nasal epithelial cells. (B) Representative short circuit current (Isc) tracings of nasal cells from a control. Amiloride (100μM) was added to inhibit epithelial sodium channels, forskolin (10μM) and IBMX (100μM) were added to activate CFTR, DMSO (0.1%, vehicle control for drug delivery) was added to compare with drug tested samples, and Inh-172 (10μM) was added to inhibit CFTR. (C) Bar chart of CFTR function in nasal cells from 24 controls. Each dot represents one recording; replicate recordings ranged from 1 to 10 per sample. (D) Histogram of CFTR function in controls. The data are normally distributed (D’Agostino and Pearson test, Anderson-Darling test, Shapiro-Wilk test, and Kolmogorov-Smirnov test). (E) Plot of mean and SD of CFTR function in male and female control samples. (Unpaired t test). (F) Plot of mean and SD of CFTR function in control samples divided into three age groups. *p≤0.05 (Ordinary one-way ANOVA with Tukey’s multiple comparisons post hoc test).

To establish the average level of CFTR function in controls, we obtained HNEs from 24 individuals with two functioning *CFTR* alleles and no pulmonary symptoms. Absence of deleterious variants was confirmed by sequencing of the whole *CFTR* gene in 18 of the controls^11^ and health status was determined by verbal report or survey instrument (**see Supplemental information**). Mean age for control participants was 32.3±2.8 (mean±SEM), of which 58.3% were females and 66.7% self-described their ancestry as white (**Table 1**). CFTR function was measured in 1-10 replicates from each participant (mean±SEM: 5.1±0.5). Mean CFTR function in 24 controls was 16.1µA/cm^2^ (SEM: 1.1µA/cm^2^; range: 4.9 to 25.2µA/cm^2^; **Figure 1C**) and measurements conformed to a normal distribution (**Figure 1D**). The mean and distribution of CFTR function did not differ by sex (female mean±SEM: 15.5±1.6µA/cm^2^; male mean±SEM: 16.9±1.5µA/cm^2^; **Figure 1E**) or between participants in their 20s and 30s (**Figure 1F**). Participants 40 years and over had lower CFTR function than the younger adults (age 20-29 CFTR function mean±SEM: 18.3±1.1µA/cm^2^; age 30-39 CFTR function: 14.6±2.4µA/cm^2^; age 40+ CFTR function: 10.6±2.4µA/cm^2^), but recruitment numbers were limited for that age group (n=4). Of the controls that submitted surveys, only one reported marijuana smoking in the 30 days prior to survey completion and none reported recent cigarette, pipe tobacco, cigar, or vape use. Four of the controls were asymptomatic according to study criteria but reported having a cough or phlegm; their CFTR function did not differ significantly from the rest of the controls (n=20). Further, five individuals with no CF-causing variants and initially recruited as controls were excluded from the study because they were symptomatic, but their CFTR function was not significantly different from what was measured in the n=20 asymptomatic controls with no cough or phlegm (**Figure S4**).

**Table 1:**
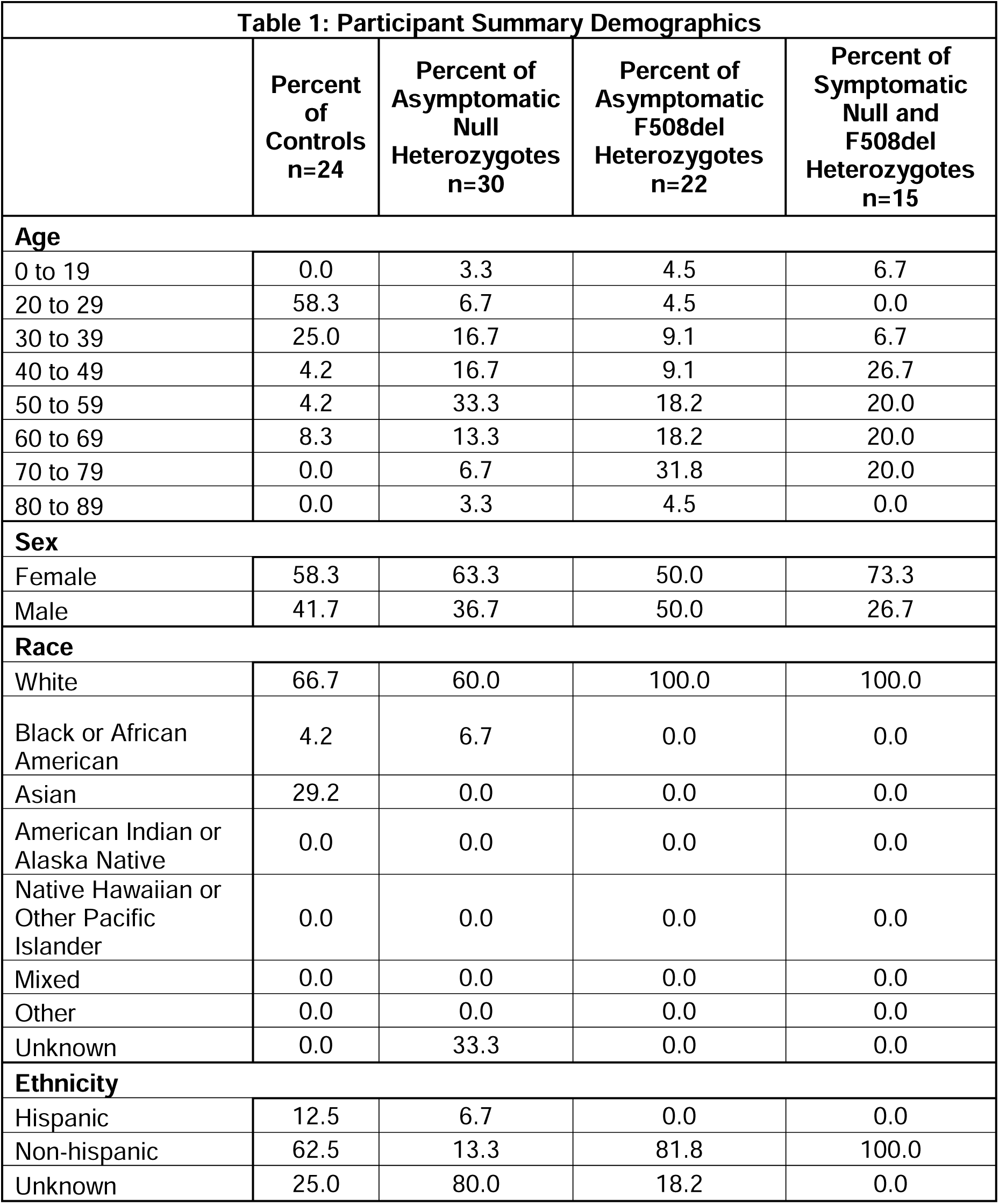
Participant Summary Demographics. Participants self-reported their age, sex, race, and ethnicity. The values in the table are percentages of the total number of people in each study group.

| **Table 1: Participant Summary Demographics** |  |  |  |  |
| --- | --- | --- | --- | --- |
|  | **Percent of Controls**<br>**n=24** | **Percent of Asymptomatic Null Heterozygotes**<br>**n=30** | **Percent of Asymptomatic F508del Heterozygotes**<br>**n=22** | **Percent of Symptomatic Null and F508del Heterozygotes**<br>**n=15** |
| **Age** |  |  |  |  |
| 0 to 19 | 0.0 | 3.3 | 4.5 | 6.7 |
| 20 to 29 | 58.3 | 6.7 | 4.5 | 0.0 |
| 30 to 39 | 25.0 | 16.7 | 9.1 | 6.7 |
| 40 to 49 | 4.2 | 16.7 | 9.1 | 26.7 |
| 50 to 59 | 4.2 | 33.3 | 18.2 | 20.0 |
| 60 to 69 | 8.3 | 13.3 | 18.2 | 20.0 |
| 70 to 79 | 0.0 | 6.7 | 31.8 | 20.0 |
| 80 to 89 | 0.0 | 3.3 | 4.5 | 0.0 |
| **Sex** |  |  |  |  |
| Female | 58.3 | 63.3 | 50.0 | 73.3 |
| Male | 41.7 | 36.7 | 50.0 | 26.7 |
| **Race** |  |  |  |  |
| White | 66.7 | 60.0 | 100.0 | 100.0 |
| Black or African American | 4.2 | 6.7 | 0.0 | 0.0 |
| Asian | 29.2 | 0.0 | 0.0 | 0.0 |
| American Indian or Alaska Native | 0.0 | 0.0 | 0.0 | 0.0 |
| Native Hawaiian or Other Pacific Islander | 0.0 | 0.0 | 0.0 | 0.0 |
| Mixed | 0.0 | 0.0 | 0.0 | 0.0 |
| Other | 0.0 | 0.0 | 0.0 | 0.0 |
| Unknown | 0.0 | 33.3 | 0.0 | 0.0 |
| **Ethnicity** |  |  |  |  |
| Hispanic | 12.5 | 6.7 | 0.0 | 0.0 |
| Non-hispanic | 62.5 | 13.3 | 81.8 | 100.0 |
| Unknown | 25.0 | 80.0 | 18.2 | 0.0 |

### HNEs from asymptomatic null and F508del CF heterozygotes generate 50% of the CFTR function measured in controls

To establish the level of CFTR function in asymptomatic CF heterozygotes and to independently replicate results, we recruited two groups of individuals; those carrying a ‘null’ variant that causes complete loss of CFTR function and those carrying the common F508del variant that decreases CFTR function to <2% of WT.^12^ The presence of a null or F508del variant in one allele and the absence of a deleterious variant in the other allele was confirmed by sequencing of the whole *CFTR* gene in 21 of the 52 asymptomatic heterozygotes (**Table S2**). In 30 null CF heterozygotes, the presence or absence of CF-like respiratory symptoms was determined by verbal report at the time of sampling (n=25) or by responses to the health history survey (n=5). Multiple replicates were measured per CF null heterozygote sample (mean±SEM; 3.4±0.2; min: 2; max: 6). Mean CFTR function was 8.8µA/cm^2^ (SEM: 0.5µA/cm^2^; min: 4.5µA/cm^2^; max: 18.7µA/cm^2^; range: 14.2µA/cm^2^; **Figure 2A**) and the distribution of individual measurements did not conform to a normal distribution due to the presence of two high function outliers (**Figure 2B**).

**Figure 2:**
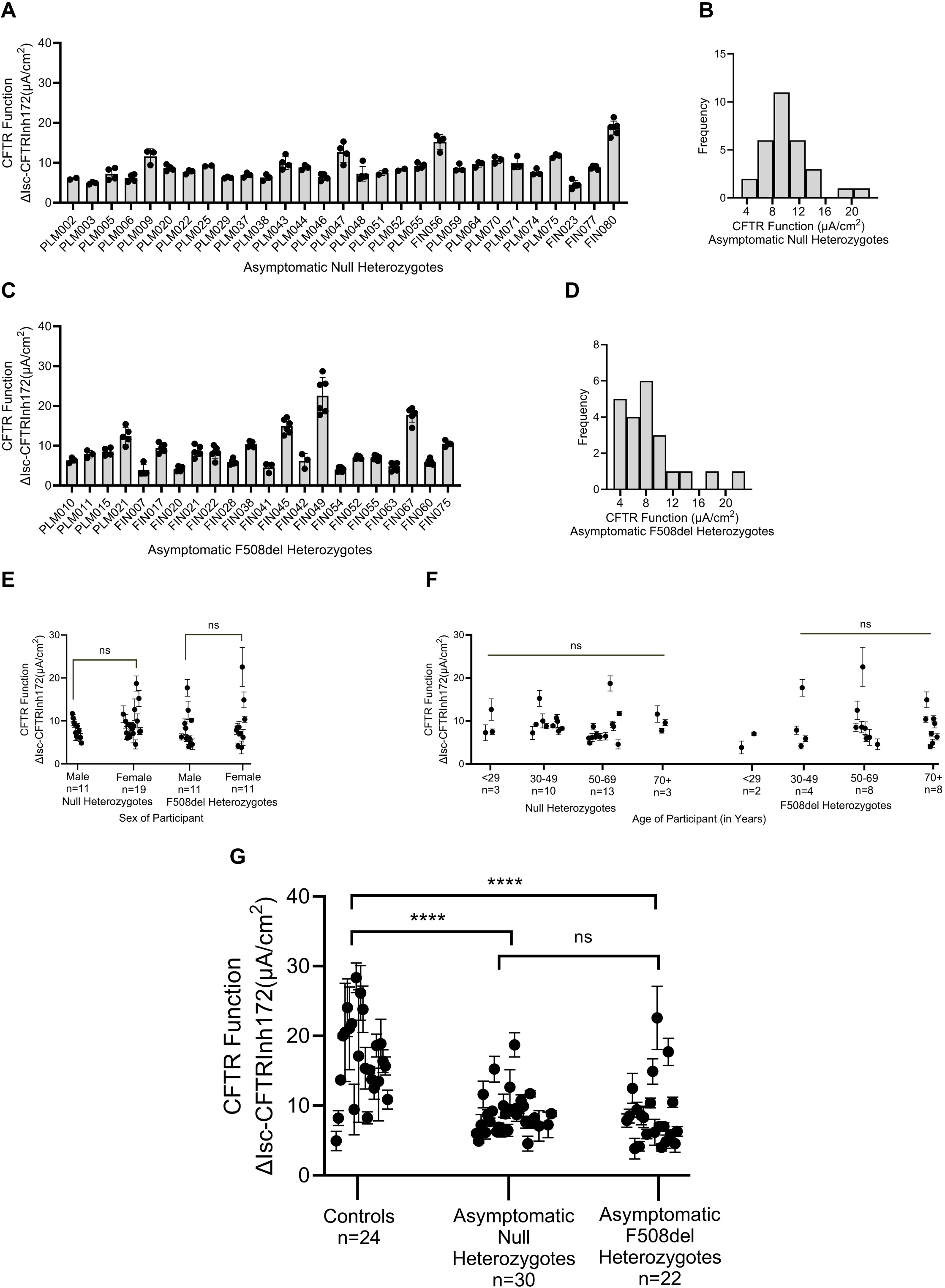
CFTR function in primary nasal cells from asymptomatic CF heterozygotes compared to controls. (A) Bar chart of CFTR function in n=30 asymptomatic null heterozygotes. Each dot represents one recording; replicate recordings ranged from 2 to 6 per sample. (B) Histogram of CFTR function in asymptomatic null heterozygous samples. (C) Bar chart of CFTR function in n=22 asymptomatic F508del heterozygotes. Each dot represents one recording; replicate recordings ranged from 3 to 6 per sample. (D) Histogram of CFTR function in asymptomatic F508del heterozygous samples. (E) Plot of mean and SD of CFTR function in male and female asymptomatic CF heterozygous samples. (Mann-Whitney test). (F) Plot of mean and SD of CFTR function in asymptomatic CF heterozygous samples divided into 4 age groups. (Kruskal-Wallis test). (G) Plot of mean and SD of aggregate CFTR function data in controls, asymptomatic null CF heterozygotes, and asymptomatic F508del CF heterozygotes. ****p≤0.0001 (Kruskal-Wallis test with Dunn’s multiple comparisons post hoc test). Normality of data distribution was assessed by D’Agostino and Pearson, Anderson-Darling, Shapiro-Wilk, and Kolmogorov-Smirnov tests.

Variability in CFTR function was also observed in samples from 22 asymptomatic F508del heterozygotes whose health status was confirmed in the same manner (verbal n=4; survey n=18). Between 3 and 6 CFTR function measurements were completed for each F508del heterozygote (mean±SD: 4.8±0.3). Mean CFTR function was 8.7µA/cm^2^ (SEM: 1.0µA/cm^2^; min: 3.8µA/cm^2^; max: 22.6µA/cm^2^; range: 18.7µA/cm^2^; **Figure 2C**) and the measurements did not conform to a normal distribution **(Figure 2D**).

For both groups of heterozygotes, CFTR current did not differ by sex (**Figure 2E**) or age group (**Figure 2F**). Given that CFTR function ranges and means were similar across all age groups for heterozygotes (range 9 to 84 years), these results indicate that CFTR function does not decline as individuals age. Of the asymptomatic heterozygotes with completed surveys, two reported smoking marijuana within 30 days prior to survey completion, one reported vaping the week before survey completion, and none reported recent cigarette, pipe tobacco, or cigar use. Despite biological variability, mean CFTR function in the asymptomatic F508del heterozygotes (8.7µA/cm^2^) was remarkably similar to the null heterozygotes (8.8µA/cm^2^). Notably, CFTR function in both groups of heterozygotes was significantly different from controls (**Figure 2G**); mean CFTR function in each group was approximately half (54.6% WT for null heterozygotes and 53.9% WT for F508del heterozygotes) of the CFTR function measured in controls.

The observation that CF heterozygotes have ∼50% of the CFTR function of controls was further substantiated through a secondary cell-based model system. Immortalized CF bronchial epithelial cells (CFBE41o-) were modified to express two *CFTR* copies from a single integrated DNA construct. F508del heterozygous cells generated currents with a mean of 47.0±7.2% WT. Information about the generation of this model and results can be found in the **Supplemental text and Figures S5, S6**.

### Airway epithelial cell type distribution does not differ between controls and CF heterozygotes

Based on observations in genetically modified ferrets and genetically corrected primary airway cells that functional CFTR is trophic for the differentiation of airway cells,^13,14^ we wanted to determine if a ∼50% reduction in CFTR function in CF heterozygotes affected the distribution of cells in HNE cultures. To address this question, we performed single cell RNA sequencing of nasal cell samples from two controls and three c.3845G>A (W1282X) null CF heterozygotes. Cell types were assigned by cell clustering followed by visualization of cell specific marker genes within an integrated dataset of all samples together (**Figure 3A, S7**). Overall cell distributions in both controls and CF heterozygotes were similar to that previously reported in direct and cultured nasal cells.^15–17^ Comparison of cell type proportions between the controls and null heterozygotes revealed no statistically significant differences, even among secretory cells and ionocytes, which are the primary sites of *CFTR* expression (**Figure 3B**).

**Figure 3:**
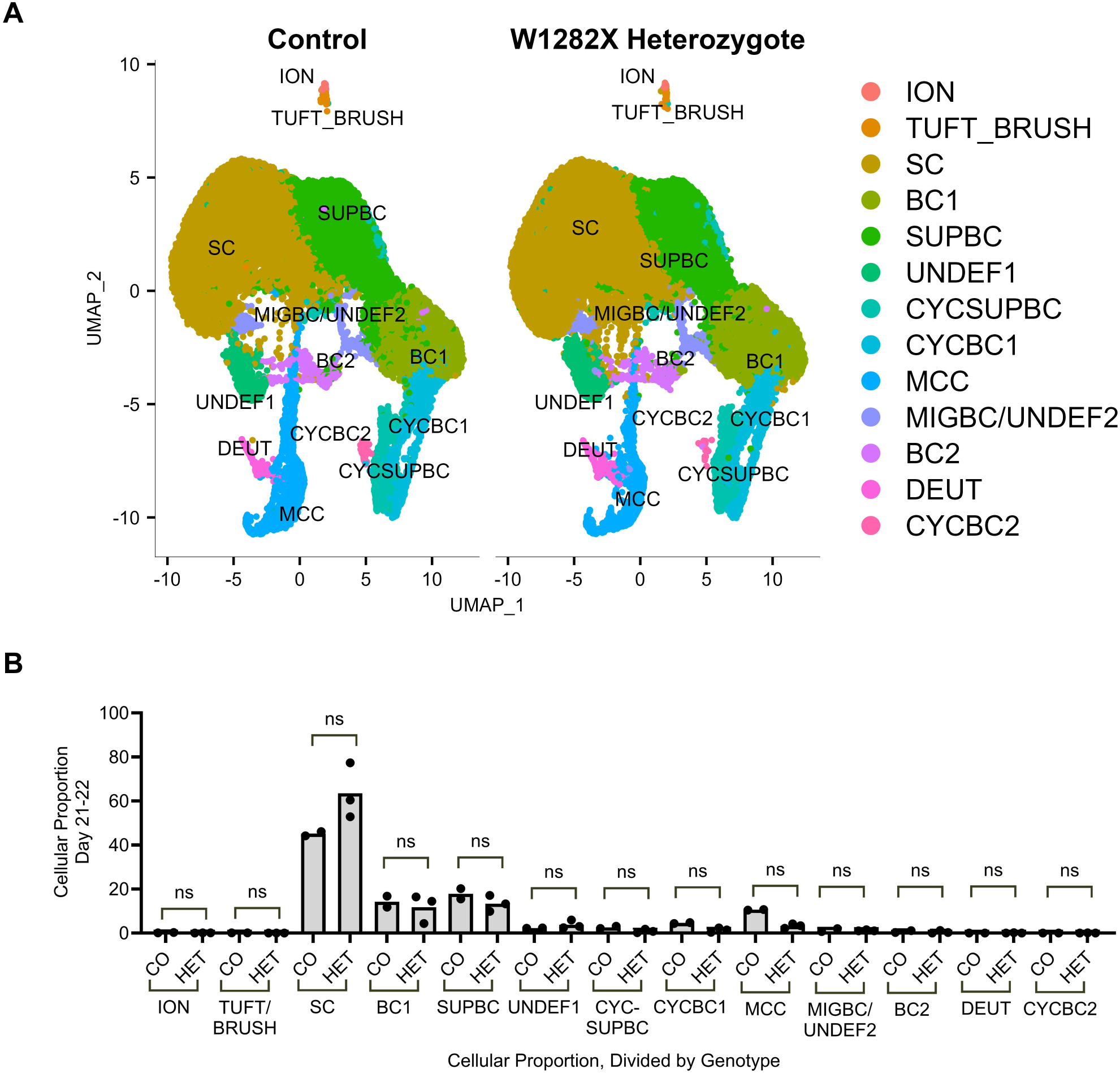
Cell type distribution in control and CF heterozygous samples. Single cell RNA sequencing was performed on nasal epithelial cells from 2 controls and 3 null heterozygous samples that had been differentiated for 21-22 days. (A) Dimensional reduction of all cell types, visualized by UMAP. (B) Plot of cell proportions in control (CO) and c.3845G>A (W1282X) null heterozygous (HET) samples. (Mann-Whitney test; ns = not significant). ION = ionocytes; TUFT_BRUSH = tuft/brush cells; SC = secretory cells; SUPBC = suprabasal cells; UNDEF1 = undefined cell cluster 1; MIGBC/UNDEF2 = migratory basal cells and undefined cluster 2; BC1 = basal cell cluster 1; BC2 = basal cell cluster 2; DEUT = deuterosomal cells; MCC = multiciliated cells; CYCBC1 = cycling basal cell cluster 1; CYCBC2 = cycling basal cell cluster 2; CYCSUPBC = cycling suprabasal cells

### CFTR function in HNEs does not differ between symptomatic and asymptomatic heterozygotes

To determine whether the presence of symptoms in some CF heterozygotes could be attributed to a reduction in CFTR function, we compared CFTR function in symptomatic and asymptomatic heterozygotes. The presence of F508del-*CFTR* in one allele and the absence of a deleterious variant in the other allele was confirmed by sequencing the whole *CFTR* gene for 14 of the 15 symptomatic heterozygotes (**Table S2**), and the presence of CF-like symptoms was determined by responses to the health history survey (15 of 15 individuals). Multiple replicates were measured per sample (mean±SEM; 4.3±0.4; min: 2; max: 6). Mean CFTR function was 8.4µA/cm^2^ (SEM: 1.0µA/cm^2^; min: 2.9µA/cm^2^; max: 18.8µA/cm^2^; range: 15.9µA/cm^2^; **Figure 4A**) and the data passed three of four normality tests (**Figure 4B**). CFTR current did not differ by sex (**Figure 4C**) or age group (**Figure 4D**). Only two of the symptomatic heterozygotes reported smoking and/or vaping marijuana within the 30 days prior to survey completion, and none reported recent cigarette, pipe tobacco, or cigar use. Notably, there was no statistically significant difference in CFTR function between the symptomatic and asymptomatic heterozygotes (**Figure 4E**). We conclude that both asymptomatic and symptomatic heterozygotes have about half of the CFTR function measured in controls (54.3% for asymptomatic heterozygotes and 52.1% for symptomatic heterozygotes).

**Figure 4:**
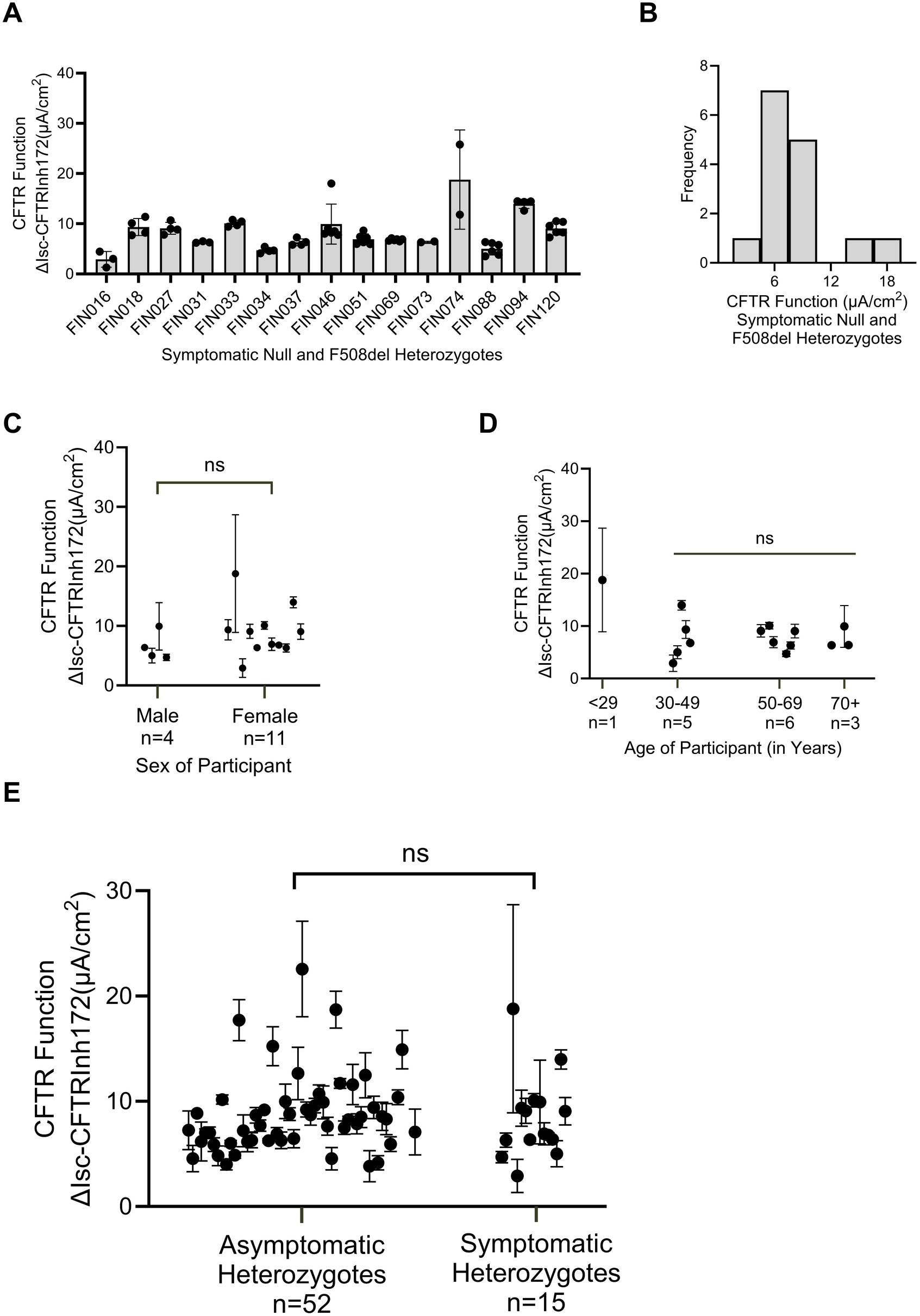
CFTR function does not differ between symptomatic and asymptomatic heterozygotes. (A) Bar chart of CFTR function in n=15 symptomatic F508del heterozygotes; 2-6 replicates were measured per sample. (B) Histogram of CFTR function in symptomatic heterozygous samples; data passed three of four normality tests (D’Agostino and Pearson test, Anderson-Darling test, Shapiro-Wilk test, and Kolmogorov-Smirnov test). (C) Dot plot (mean and SD) comparing CFTR function in male and female symptomatic heterozygous participants. (Unpaired t test). (D) Dot plot (mean and SD) comparing CFTR function in symptomatic heterozygous samples of varying ages. (Kruskal-Wallis test). (E) Dot plot (mean and SD) comparing aggregate CFTR function data in asymptomatic and symptomatic heterozygotes having one null or F508del-*CFTR* allele. (Mann-Whitney test).

### Response to CFTR modulators is variable between symptomatic CF heterozygotes

To determine whether CFTR modulators could be a viable treatment option for symptomatic CF heterozygotes, we treated primary nasal cultures with the CFTR modulators ivacaftor and elexacaftor-tezacaftor-ivacaftor (ETI) to determine whether CFTR function could be increased above baseline (i.e. no CFTR modulator drugs). Nasal cells from ten of the symptomatic heterozygotes and one of the asymptomatic null heterozygotes were subjected to modulator testing. We measured several replicates per condition (baseline: mean±SEM 3.7±0.3; min: 2; max: 6; ivacaftor: mean±SEM 2.8±0.2; min: 2; max: 4; ETI: mean±SEM 3.4±0.2; min: 2; max: 4). Statistics were performed on samples with greater than two replicates, and response to CFTR modulators was variable across samples (**Figure 5**). Four of eleven samples exhibited increased CFTR function when exposed to ivacaftor and seven of eleven samples exhibited a significant increase in CFTR function with ETI treatment compared to baseline. Notably, sample FIN016 had lower baseline CFTR function than all other samples subjected to modulator testing and treatment with ETI increased CFTR function to WT levels. Interestingly, two samples had no response to any modulators: ivacaftor alone (FIN027) or ETI (FIN035, FIN027). Two other samples with only two replicates (FIN073 and FIN074) also appeared unresponsive to modulators. We conclude from these studies that CFTR modulators may be a beneficial treatment option for some CF heterozygotes with symptoms but not for others.

**Figure 5:**
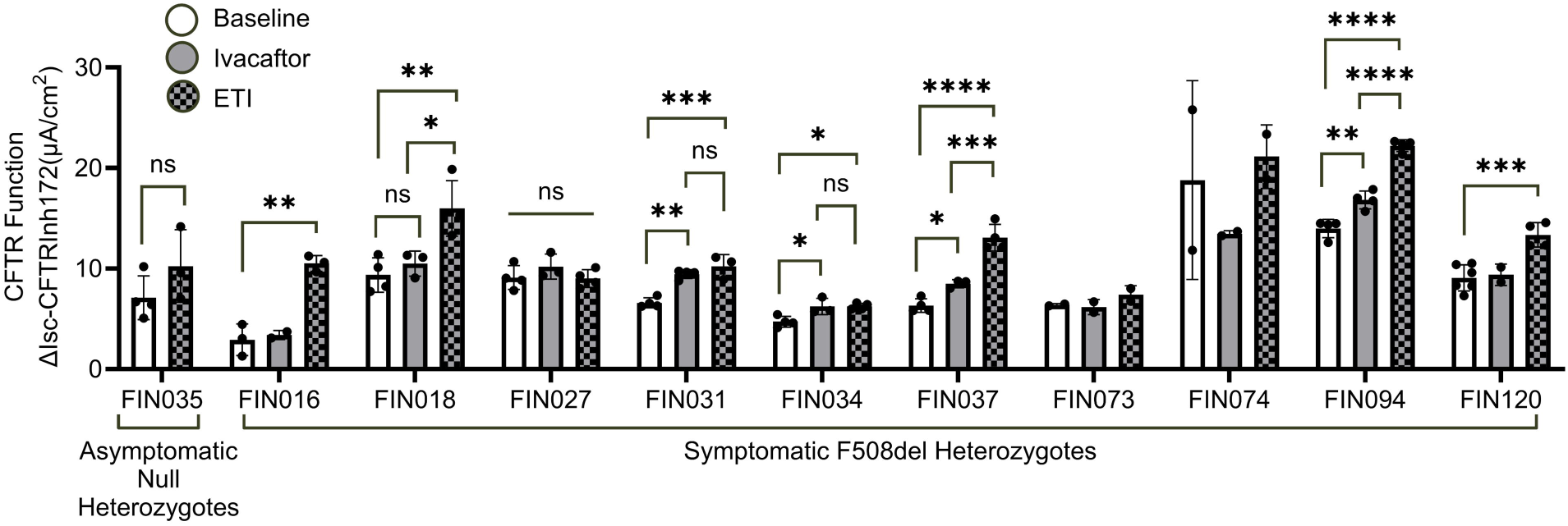
Primary nasal cells exhibit varied response to CF modulators. CFTR function was measured in nasal cells from 11 heterozygous participants under 3 conditions: baseline (no modulators, DMSO only), ivacaftor only, or ETI (elexacaftor-tezacaftor-ivacaftor). Bar chart of CFTR function per sample and per condition. Dots represent individual measurements. Statistical comparisons were performed for samples with more than 2 replicates per group; thus, no statistical comparisons are shown for FIN073 and FIN074, which had fewer than 3 measurements per condition. Groups with more than two data points were subjected to normality tests; all groups passed the Shapiro-Wilk test except FIN031 ivacaftor and FIN094 baseline. Two group comparisons: Unpaired t test. Three group comparisons: Ordinary one-way Anova with Tukey’s multiple comparisons post hoc test.

## Discussion

In this study we addressed two questions: what is the level of CFTR function in CF heterozygotes and is CFTR function lower in symptomatic CF heterozygotes than asymptomatic heterozygotes? This subject is relevant because CF heterozygotes have been reported to be at increased risk for a number of CF-related respiratory disorders (**Supplemental Table S1**) and there are estimated to be 25 million CF heterozygotes worldwide. We demonstrate that mean CFTR function in HNEs from CF heterozygotes carrying a null variant or F508del is approximately 50% of the mean recorded in controls, consistent with the expectation of a recessive Mendelian disorder. However, the considerable variability we observed within study groups indicates that the level of CFTR function sufficient for normal lung function differs among people. This supposition is supported by our observation that respiratory symptoms in CF heterozygotes do not correlate with a reduction in CFTR function and by the observation that controls with respiratory symptoms do not have reduced CFTR function. Our results suggest that reduction in CFTR function predisposes heterozygotes to respiratory disease, but individual factors (i.e., environmental exposures, modifier genes) are responsible for the appearance of CF-like symptoms. The important role of CFTR-independent factors to the evolution of CF lung disease has been documented by studies of CF twins and siblings.^18,19^ In sum, our results indicate that a single functional CFTR gene is sufficient to avoid the development of CF, but in a fraction of individuals, it is insufficient to prevent the development of respiratory symptoms when certain exposures or genetic variants are present.

Previous studies of CFTR function in CF heterozygotes using nasal potential difference (NPD), sweat chloride concentration, and cell-based chloride transport have reported substantial variability in each measure and overlap among controls, heterozygotes, and individuals with CF. In a study of NPD in 37 CF heterozygotes and 29 control participants, almost half (n=17; 46%) of the CF heterozygotes had CFTR chloride transport that was greater than the mean in controls.^20^ However, when considered as a group, the mean function in heterozygotes was about half that observed in controls (-8.2 vs -15; p =0.0002). The authors were unable to conclude whether CF heterozygotes had half of the CFTR function in controls. Another NPD study revealed no statistically significant difference in the sodium and chloride diffusion potential between 25 controls and 21 CF heterozygotes.^21^ Measurements of sweat chloride concentration have shown significant differences in means between individuals with CF and heterozygotes, but not between controls and heterozygotes.^20,21^ Cell-based assessments of CFTR function in airway epithelial cells using SPQ (6-methoxy-\(N\)-(3-sulfopropyl)quinolinium) fluorescence of individual nasal epithelial cells also yielded inconsistent results. One third of 17 heterozygotes had values overlapping with 77 individuals with CF, and half of the heterozygotes had values less than 50% of that seen in controls.^20^ Recently, a small-scale study of human bronchial epithelial cells from two CF heterozygotes reported CFTR function to be ∼80% of wild type CFTR function.^22^ While these studies provided evidence that CFTR function in heterozygotes differs from controls, inconsistency due to broad technical variability in NPD^23–25^ and sweat chloride measurement^26–28^ as well as the relatively small size of the cell-based studies precluded a quantitative determination. Recognizing the challenges of technical and biologic source of variability, we elected to use a robust primary cell assay to obtain a reliable measure of CFTR function in a sufficient number of subjects to enable comparison of group means. Replication of results from null heterozygotes in a group of F508del heterozygotes and in a model cell system increases our confidence that individuals with one functioning allele have, on average, 50% of the CFTR function of individuals with two functioning alleles.

Our observations have several implications for treatment of CF. First, mixing studies have shown that blending CF cells with 5-20% of cells corrected to ‘wild type’ levels of CFTR function corrected chloride transport to 70% to 100% of normal.^29–32^ These studies imply that correcting CFTR function in a small fraction of cells will be therapeutic. However, cell monolayers grown from nasal epithelia of CF heterozygotes with one functioning WT-*CFTR* allele per cell generate about 50% of the chloride transport observed in monolayers of individuals with two functioning CFTR alleles. We obtained the same result when we used a model cell line expressing one or two CFTR alleles, and we excluded differences of expression in cell subtypes between heterozygotes and controls using single cell RNA sequencing. Our result suggests that gene correction may have to be pervasive in airway epithelia, as is achieved by small molecule CFTR modulators. Second, given the range of CFTR function measurements we recorded in HNEs from controls and asymptomatic CF heterozygotes, the level of CFTR transport in airway epithelia required to escape CF symptoms differs among individuals. Application of the same threshold for participants in a clinical trial may fail to detect individuals who have therapeutic increases in CFTR function despite falling short of a single target. Testing baseline and post treatment CFTR function in epithelial cell monolayers could allow individualized precision treatment based on gene correction methods. Third, biochemical studies have shown that CFTR may dimerize under certain conditions,^33^ and that F508del-CFTR protein can affect the function of “wild-type” CFTR produced by the second CFTR allele.^34^ If such interactions truly occur, we would expect that the function of “wild-type” CFTR would be compromised in a heterozygote. However, we noted that HNEs from F508del heterozygotes had the same level of CFTR function than cells from null heterozygotes; and function in both groups of heterozygotes was about half that recorded in individuals with two working *CFTR* alleles. While our observations do not exclude the possibility that a different *CFTR* variant may produce a protein that interferes with the product from the second *CFTR* allele, we present evidence that F508del-CFTR does not interfere with the “wild-type” *CFTR* protein. Fourth, the lack of difference in CFTR function between asymptomatic and symptomatic CF heterozygotes indicated that augmentation with CFTR modulators is unlikely to be clinically effective. The uncertain clinical effectiveness of increasing CFTR function is emphasized by the variable response of HNEs from CF heterozygotes to CFTR modulators.

There are several limitations to this study. We defined symptomatic individuals using only respiratory symptoms but CF-like symptoms in other organ systems have been reported at increased prevalence in CF heterozygotes.^35^ On the other hand, replication of the increased risk of intestinal and pancreatic complications in CF heterozygotes is lacking, whereas disease of the sinuses and airways have been consistently reported at higher prevalence in heterozygotes.^36,37^ The age distribution differed between controls and CF heterozygotes; however, we presented evidence in CF heterozygotes that CFTR function does not decrease as people age. The system we used for measuring CFTR function only accounts for CFTR chloride transport and does not account for other reported CFTR functions, such as bicarbonate secretion, mucus transport or epithelial sodium channel regulation.^38–40^ Other functions of CFTR might be disrupted to a different degree in CF heterozygotes, especially in the symptomatic heterozygotes even though chloride transport has been the CFTR function consistently correlated with CF severity.^41^ Unlike controls, CFTR function in the asymptomatic null and F508del CF heterozygotes did not conform to a normal distribution. This distortion appears to be caused by outliers in which the high amount of CFTR function measured may be due to some unresolved technical artifact or an unrecognized biological or environmental variable. Longitudinal sampling from the same subjects in a future study could better address reproducibility and possibly factors that contribute to individual variability in CFTR function, as shown for studies of CFTR modulator effect.^42^

## Conclusion

The increased use of genetic testing has dramatically increased the number of individuals aware of their CF heterozygous status. Given that there are an estimated 25 million CF heterozygotes worldwide, it is important that physicians understand how to counsel these individuals regarding their risk for chronic respiratory diseases and treatment options. CF heterozygotes have, *on average*, 50% of the amount of CFTR function seen in controls, and there is no correlation between CFTR function and symptoms in heterozygotes. The variability within study groups indicates that individuals have unique CFTR functional thresholds required to avoid developing CF-like symptoms and other factors (i.e. genetic, environmental, and stochastic) should be considered as contributors to symptom development

## Supporting information

Health History Survey

Chronic Bronchitis Flow Chart

Chronic Rhinosinusitis Flow Chart

Recurrent Acute Rhinosinusitis Flow Chart

## Authors’ Contributions

Conceptualization – G.C., A.B.

Data curation – A. Pion, A.B., N.S.

Formal analysis – A. Pion, A.B., A.E., K.R., C.M., N.S.

Funding acquisition – A.B., G.C.

Investigation – A. Pion, A.B., E.K., A.T., K.S., A. Parr, N.S.

Methodology – A.B., G.C.

Project administration – A. Pion, A.B., N.S., G.C.

Resources – P.M., C.M., N.S., G.C.

Software – A. Pion, A.E., K.R., C.M.

Supervision – G.C.

Validation – A. Pion, A.B., G.C.

Visualization – A. Pion, A.B., G.C.

Writing – original draft – A. Pion, G.C.

Writing – review & editing – A. Pion, G.C.

## Conflicts of interest section

K.R. is a consultant for BillionToOne and Vertex Pharmaceuticals (Europe) Limited.

## Funding section

This work was funded by NIDDK R01 DK44003 and a grant from the Spruance Foundation awarded to G.C. N.S. was supported by a Research Innovation Award from Vertex Pharmaceuticals and by Cystic Fibrosis Foundation grant SHARMA23GO. A.P. was funded in part by a grant from NIGMS T32GM148383.

## Acknowledgements section

The authors would like to thank all participants for making this project possible. We would like to thank Molly Sheridan and David Mohr for their assistance with DNA preparation, sequencing, and analyses. We would also like to thank all the physicians and staff members that aided us in our recruitment efforts, especially Donna Peeler, Kaia Houtman, Shruti Paranjape, Elisa Ignatius, Tia Day, Natalie West, Lori Vanscoy, Jamelia Maynard, and Peter Mogayzel.

## Data availability section

Single cell RNA sequencing data will be uploaded to GEO after paper submission.

## Statement of research impact on clinical medicine

CF heterozygotes are at increased risk for CF-like symptoms and there are estimated to be 25 million CF heterozygotes worldwide; taken together, this presents a significant disease burden that should be addressed. The work presented in this manuscript informs patient-facing personnel on how to advise CF heterozygotes about their lifetime disease risk, underlying disease cause, and treatment.

