## Supplemental Text for "CFTR function in nasal airway cells from symptomatic and asymptomatic CF heterozygotes"

**Title of submission:** **CFTR function in cystic fibrosis heterozygotes**

**Authors’ names:**

**Participant recruitment**
Participants were recruited through posters in Johns Hopkins University (JHU) buildings and email notifications sent to JHU staff. CF heterozygotes were specifically recruited from the JHU Cystic Fibrosis Center and JHU Adult Pulmonology clinic. Also, 555 letters were mailed to JHU CF patients addressing their CF heterozygous family members (i.e. parents, siblings, and children of patients with CF). Participants were consented under approved studies (IRB00116966 and/or IRB00235883). Heterozygous participants were excluded from the study if they harbored a CF-causing variant that was not null-*CFTR* or F508del-*CFTR*, had the inferior turbinate of both nostrils removed, or took blood thinners. A human nasal epithelial (HNE) study participant list with anonymized study identifiers, genotype, phenotype, and differentiation media regimen can be found in **Supplemental Table 2**. A pedigree table for all participants who provided HNEs can be found in **Supplemental Table 3**.

**Isolation of HNEs**
Nasal brushings were performed by trained physicians at JHU. 2% topical tetracaine was first administered to both nostrils. Cells were collected by rotating cytology brushes on the superior surface of inferior turbinates. Brushes were immediately submerged in HNE expansion media (**Supplemental Table 4**) and cultured.

**HNE Culture and Measurement of CFTR function in HNEs by short circuit current**

HNE cells were cultured and CFTR function was measured as previously described with only a few notable differences.^1^ The media constituents and their concentrations are listed in **Supplemental Table 4**. The differentiation media regimen differed for some samples: For cultures differentiated for 28 days and profiled by single cell RNA sequencing, the media regimen was PALI for 14 days, VALI 5 days, PALI 2 days, VALI 5 days, and PALI 2 days. Also, among the cultures which were the first to be collected, some were differentiated completely in VALI media. Media regimen by sample is listed in **Supplemental Table 2**. All CF modulators were purchased from Selleck Chemicals (catalog numbers: ivacaftor [S1144], elexacaftor [S8851], tezacaftor [S7059]).

**Saliva collection and gDNA isolation**
About 3mL of saliva was collected from each participant in saliva collection tubes (manufacturer: DNA Genotek; product name: Oragene^TM^ dx OGD-600). Samples were kept at room temperature or on ice after collection and refrigerated at 4^o^C until gDNA extraction. gDNA was isolated according to the manufacturer’s protocol then stored at –20^o^C.

**Full gene *CFTR* sequencing**
Full gene *CFTR* sequencing and analysis of sequencing results was performed as previously

described.^2^

**Health history survey assembly and response evaluation**

The health history survey was adapted from the CF patient registry questionnaire, the St. George’s Respiratory questionnaire, and the American Thoracic Society Division of Lung Disease questionnaire. The full survey titled “Health History Survey” is included with the supplemental materials. Participants were classified as symptomatic if any of the following were true: A) They answered “yes” to the questions “Have you ever had chronic bronchitis?” and “Was it confirmed by a doctor?” B) They answered “yes” to the question “Has a doctor ever told you that you have bronchiectasis?” or C) Comparison of their survey responses to a flow chart (included as additional materials entitled “Chronic Bronchitis Flow Chart,” “Chronic Rhinosinusitis Flow Chart,” and “Recurrent Acute Rhinosinusitis Flow Chart”) of relevant survey questions indicated that they had chronic bronchitis, chronic rhinosinusitis, or recurrent acute rhinosinusitis. To evaluate C, flow charts were curated based on the clinical definitions for chronic bronchitis, chronic rhinosinusitis, or recurrent acute rhinosinusitis. (One notable exception is that the definitions for chronic rhinosinusitis and recurrent acute rhinosinusitis require that the nasal discharge be purulent (i.e. cloudy); we believed that there was a high likelihood for participants to inaccurately recall the nature of their nasal discharge and thus disregarded the nasal purulence question.) The survey responses were reviewed independently by three individuals – one of which is a practicing pulmonologist - and the above questions and flow charts were followed strictly for phenotype classifications. Study records were scored objectively as symptomatic or asymptomatic using separately coded computer algorithms by two of the researchers who were masked to participant’s genetic and CFTR functional data; the third researcher scored symptomatology by hand for each record. Discrepancies were discussed between the researchers until a consensus was reached.

**Assessment of CFTR function in cell lines expressing one or two WT CFTR cDNAs**

Generation of Dual Expressing Cell Lines:

The ThermoFisher Scientific kpEF5/FRT/V5-D-TOPO plasmid was genetically modified through Gibson Assembly to express *CFTR* from two separate *CFTR* alleles, with each allele having its own EF-1alpha promoter and poly(A) signal and alleles were separated by two copies of the chicken hypersensitive site 4 insulator (cHS4; **Figure S5A**). One plasmid had two wild type *CFTR* alleles (WT/WT), one plasmid had one wild type and one F508del-*CFTR* allele (WT/F508del), and one plasmid had two F508del-*CFTR* alleles (F508del/F508del). To differentiate between alleles, three base pairs in positions 79-81 of the 3’ UTR of one *CFTR* allele were deleted by site directed mutagenesis, and the two alleles also differed at the c.1408 location; c.1408A (M470) in one allele, c.1408G (V470) in the other. Plasmid authenticity was verified by Sanger sequencing. Independent transcription of each *CFTR* cDNA was tested by transient transfection of WT/WT plasmid into HEK293 cells, RNA extraction, and RT-PCR using three primer sets bridging the 3’ end of one allele and the 5’ end of the second allele. Amplified products were separated and sized using gel electrophoresis and compared to amplification of the plasmid DNA as a control as shown in **Figure S5B**. CF bronchial epithelial (CFBE41o-; abbreviated CFBE) cells with a Flp-In recombination target site were transfected with one of the three designed plasmids and cell clones with integrated plasmid were selected based on Hygromycin resistance, as previously described.^3^ Nine WT/WT-*CFTR*, four WT/F508del-*CFTR*, and four F508del/F508del-*CFTR* stable CFBE lines were made. qRT-PCR was utilized with SsoAdvanced Universal SYBR Green mix (Bio-Rad) to measure the amount of *CFTR* RNA in each CFBE clone.

Functional Testing in CFBEs:

CFBEs were grown on Snapwell^TM^ filters to a confluent monolayer. When transepithelial resistance reached ~200µΩ, filters were mounted in Ussing chambers (Physiologic Instruments) and bathed in asymmetrical buffer (higher chloride concentration on the basal side than the apical side). Voltage was clamped at zero for measurement of short circuit current. Cells were allowed time to equilibrate in the buffer before addition of experimental drugs. 10μM forskolin was added to the basal side of cells to increase CFTR function. After current stabilization, 10μM CFTR inhibitor 172 (Inh-172) was added on the apical side to inhibit CFTR function. The amount of CFTR function in a sample was calculated as the amount of short circuit current before Inh-172 addition minus the amount of short circuit current after Inh-172 addition. More details of the experimental method can be found in previous publications from our laboratory.^4,5^

Calculating CFTR function in CFBE clones as a percentage of WT/WT-*CFTR* expressing clones

The amount of CFTR function in a clone was calculated relative to WT/WT-*CFTR* expressing clones by a multi-step calculation, as described previously.^4^ In brief, the amount of *CFTR* mRNA in WT/WT-*CFTR* expressing clones was plotted versus the amount of CFTR function measured in WT/WT-*CFTR* expressing clones and a line of best fit passing through the origin was derived to find the slope (192.12). For subsequent calculations, %WT/WT was computed with the following formula:

$\%WT/WT CFTR function=\frac{\left( 100 \right)*\left( amount of function measured in clone \right)}{(192.12)(amount of RNA measured in clone)}$

**Single cell RNA sequencing**

After 14, 21, or 28 days of differentiation, a single cell suspension was made by dissociating HNEs from filters. Libraries were generated with a targeted cell recovery of 8000 cells using the 10x Genomics Chromium NextGEM Single Cell 3’ v3.1 platform following the manufacturer’s instructions. Libraries were sequenced at the JHU Genetic Resources Core Facility using the NovaSeq SP sequencer. Reads were aligned to hg38 with CellRanger, a Seurat object was generated in Seurat_4.1.0, and downstream analyses were performed in Seurat version 5.3.0 using R version 4.5.1 in RStudio version 2025.9.0.387. A total of 83,391 cells were profiled from 15 samples and datasets were individually filtered (see **Supplemental Table 5**). Samples were integrated and downstream analyses were performed on the single integrated dataset. 21 dimensions were used to find cell neighbors, unsupervised clustering was performed at a resolution of 0.8. In the UMAP, 21 PCA dimensions were reduced. To name cell clusters, marker genes were visualized using violin plots, dot plots, and feature plots within Seurat. The marker genes used were basal cells (KRT5+, TP63+ high); secretory cells (MSMB+, SCGB3A1+); multiciliated cells (FOXJ1+, DNAH5+); deuterosomal cells (FOXJ1+, DEUP1+, FOXN4+); cycling basal cells (MKI67+, KRT5+); suprabasal cells (KRT5+, TP63 low); migratory basal cells (FN1+, VIM+); and rare cells (*CFTR*+, ASCL3+).^6,7^ The ionocytes and tuft/brush cells clustered together in one group of cells which we called the “rare” cells. To determine exactly how many ionocytes versus tuft/brush cells were present, we subset the rare cells from the rest of the data, performed clustering to visualize subclusters, then analyzed marker gene expression within the subclusters (ionocytes: *CFTR*+ and FOXI1+; tuft/brush: ASCL2+ and LRMP+).^6^ The expression of select marker genes can be found in **Figure S7.** Cell clusters that did not fall into one of the above categories were labeled as “UNDEF,” which stands for “undefined.” One cluster had some migratory basal cells and other cells which were undefined; hence, we named that cluster MIGBC/UNDEF2 (for migratory basal cells / undefined).

**Graphing and statistical analyses**
All graphing and statistical analyses were performed in GraphPad prism (GraphPad by Dotmatics). Normality was assessed with the following four tests: D’Agostino and Pearson, Anderson-Darling, Shapiro-Wilk, and Kolmogorov-Smirnov. Parametric statistical tests were utilized when all groups were normally distributed and non-parametric statistical tests were utilized if one or several groups were not normally distributed. (A single exception is Figure 5: parametric tests were utilized for all comparisons regardless of whether groups were normally distributed or not to maintain consistency.) To compare two groups of data, either the unpaired t-test (parametric) or the Mann-Whitney test (non-parametric) was applied. To compare three groups of data, either the ordinary one-way ANOVA with Tukey’s multiple comparisons post hoc test (parametric) or the Kruskal-Wallis test with Dunn’s multiple comparisons post hoc test (non-parametric) was applied. For all tests, the significance threshold was p=0.05. Statistical analyses were performed on groups having at least two data points with the only exception being the comparison of cellular proportions between control and heterozygous single cell RNA sequencing data. For the single cell RNA sequencing data, the non-parametric Mann-Whitney test was used for all comparisons because of the small sample sizes.

**Graphical illustrations**

Graphical illustrations (**Figure 1A**, **Supplemental Figure 6A, Chronic Bronchitis Flow Chart, Chronic Rhinosinusitis Flow Chart, Recurrent Acute Rhinosinusitis Flow Chart**) were created using BioRender (<https://biorender.com/>).

**Legend Text for Supplemental Tables:**

**Supplementary Table 1:** **Original Research Articles About Symptoms in CF Heterozygotes** The table represents a collection of relevant research articles in which the relationship between CF heterozygous status and CF-like symptom development was explored.

**Supplementary Table 2:** **De-identified Nasal Cell Participant List** This table lists participants that contributed a nasal brushing sample and provides information about the genetic analyses performed on their sample, phenotype, CFTR functional results, and differentiation media regimen used. PALI_VALI_PALI = represents cells exposed to PALI for 14 days, VALI for 8-10 days, then PALI for 2 days before functional assessment. VALI only = cells exposed to VALI media for the entire time of cell differentiation.

**Supplementary Table 3:** **Nasal Sample Pedigree Table** This table provides information about how the participants providing nasal samples are related to each other. The “Family” column: One number is assigned per family. Thus, related individuals have the same family number. Individuals with different family numbers are unrelated. The “Individual” column: Each person within a family is assigned their own number, starting at 1. The “Father” and Mother” columns: Uses the numbers assigned in the “Individual” column to indicate who is the mother or father in the family. If a person has zeroes (“0”) entered into both the “Father” and “Mother” slots, then their parents are not included in the study. The “Sex” column uses 1 to represent males and 2 to represent females. Study identifiers ending in a letter represent “filler” slots which were necessary for pedigree table completion but do not represent real participants in the study. Conversely, study identifiers ending in a number represent study participants.

**Supplementary Table 4:** **Media Constituents** This table describes the reagents included in the three types of media utilized to culture HNEs and their concentrations.

**Supplementary Table 5: Filtering Thresholds for Single Cell RNA Sequencing Datasets** This table lists the filtering thresholds selected individually for each of the 15 single cell RNA sequencing datasets for quality control. These thresholds were utilized to subset the data. nFeatures stands for the number of genes. Percent.mt stands for the percentage of reads mapping to the mitochondrial genome.

**Supplemental Figure Legends:**

**Supplemental Figure 1: Technical variables that did not alter CFTR function** (A) Plot of CFTR function in samples from the left and right nostrils that were cultured separately for two controls (FIN001 and FIN002). P2, P3, P4 = passages 2, 3, and 4, respectively. (Mann-Whitney tests). (B) Plot of CFTR function of nasal cells from a healthy F508del het that were switched to PALI media on the basal side of filters on differentiation day 22. Cells were cultured for 2, 3, or 4 additional days then CFTR function was tested. (Ordinary one-way ANOVA).

**Supplemental Figure 2: Technical variable that had a variable effect on CFTR function** Plot of CFTR function in four samples that were exposed to a variable number of irradiated 3T3 fibroblasts upon initial seeding on Snapwell™ filters. The dot plot demonstrates CFTR function in filters exposed to a “High” versus “Low” number of 3T3s. “FIN001 High” = 1.94*10^6cells/well; “FIN001 Low” = 9.72*10^5cells/well; “FIN013 High” = 2.0*10^6cells/well; “FIN013 Low” = 6.9*10^5cells/well; “FIN016 High” = 2.0*10^6cells/well; “FIN016 None” = 0cells/well; “FIN075 High” = 2.0*10^6cells/well; “FIN075 Low” = 1.0*10^6cells/well. Statistical test used for comparisons in samples FIN001 and FIN013: Unpaired t test. There are no statistical comparison bars above samples FIN016 and FIN075 because there were fewer than three replicates per condition.

**Supplemental Figure 3: Technical variables that altered CFTR function** (A) Plot of CFTR function in samples cultured “Fresh” (i.e. not cryopreserved) and “Frozen” (cryopreserved). ****p≤0.0001, **p≤0.01 (Unpaired t test). (B) Plot of CFTR function tested at passages 2, 3, or 4 in nasal cells from a control. ****p≤0.0001 (Ordinary one-way ANOVA with Tukey’s multiple comparisons post hoc test). (C) Plot of CFTR function measured repeatedly in two cultures, one control (blue) and one c.2195T>G (L732X) heterozygote (black), between 20-30 days of differentiation. Plot displays mean (solid points) and standard deviation (dotted lines).

**Supplemental Figure 4: Reduced CFTR function is not necessary for respiratory symptom development** Plot of CFTR function in nasal cells from controls with no symptoms, controls with cough or phlegm (but asymptomatic according to study criteria), and controls symptomatic according to study criteria. Dots with error bars represent mean and standard deviation within each sample. CB= chronic bronchitis, CRS = chronic rhinosinusitis, RARS = recurrent acute rhinosinusitis, BR = bronchiectasis (Ordinary one-way ANOVA).

**Supplemental Figure 5: Dual expression plasmid design and confirmation that two distinct RNA transcripts are produced from the plasmid in HEK293 cells.** (A) Map of plasmid with two copies of full-length WT-*CFTR* cDNA each with EF-1alpha promoter and bovine growth hormone poly A signal. The *CFTR* cDNAs are separated by two copies of the chicken hypersensitive site 4 (cHS4) insulator. The two *CFTR* copies differed at c.1408 (adenine or guanine nucleotide; corresponding to methionine or valine at codon 470) and positions 79-81 of the 3’ UTR. (B) To determine if separate *CFTR* RNA transcripts were generated by each copy of *CFTR*, the region surrounding the cHS4 insulators was amplified with three primer sets (see top panel) then visualized by gel electrophoresis. The middle panel shows DNA fragments of the correct size when plasmid DNA from two separate WT/WT-*CFTR* clones were amplified. The bottom panel shows that fragments were not amplified from cDNA synthesized from RNA extracted from HEK293 cells transiently transfected with WT/WT-*CFTR.* No RT = no reverse transcriptase negative control. Water = no sample negative controls.

**Supplemental Figure 6: CFTR function in CF bronchial epithelial (CFBE) cells carrying two WT (wild-type) *CFTR* copies is distinguishable from CFTR function in cells with one WT and one F508del copy.** (A) Diagram of experimental procedure: CFBE41o- cells were stably transfected with a *CFTR* dual expression plasmid then CFTR function was measured. (B) Bar chart of CFTR function normalized by RNA quantity in CFBE41o- cells expressing two copies of WT-*CFTR* from a dual expression plasmid. Each dot represents one recording; replicate recordings ranged from 2 to 6 per sample. (C) Histogram of CFTR function normalized by RNA quantity in CFBE41o- cells expressing WT/WT-*CFTR* from a dual expression plasmid. The data are normally distributed (D’Agostino and Pearson test, Anderson-Darling test, Shapiro-Wilk test, and Kolmogorov-Smirnov test). (D) Plot of mean and SD of CFTR function across an average of 4.2±0.4 technical replicates for each CFBE clone (WT/WT-*CFTR* cDNA, n=9 clones; WT/F508del-*CFTR* cDNA, n=4 clones; F508del/F508del-*CFTR* cDNA, n=4 clones). *p≤0.05, ***p≤0.001 (Ordinary one-way ANOVA with Tukey’s multiple comparisons post hoc test).

**Supplemental Figure 7: Cell type identification by marker gene expression** Single cell RNA sequencing was performed on nasal epithelial cells from two controls and three c.3845G>A (W1282X) null heterozygous samples differentiated for 14, 21, and 28 days. Dot plot shows marker gene expression per cluster after cell naming. ION = ionocytes; TUFT_BRUSH = tuft/brush cells; SC = secretory cells; SUPBC = suprabasal cells; UNDEF1 = undefined cell cluster 1; MIGBC/UNDEF2 = migratory basal cells and undefined cluster 2; BC1 = basal cell cluster 1; BC2 = basal cell cluster 2; DEUT = deuterosomal cells; MCC = multiciliated cells; CYCBC1 = cycling basal cell cluster 1; CYCBC2 = cycling basal cell cluster 2; CYCSUPBC = cycling suprabasal cells.
