## Supplementary figures and images for "CFTR function in nasal airway cells from symptomatic and asymptomatic CF heterozygotes"

### Chronic Bronchitis Flow Chart

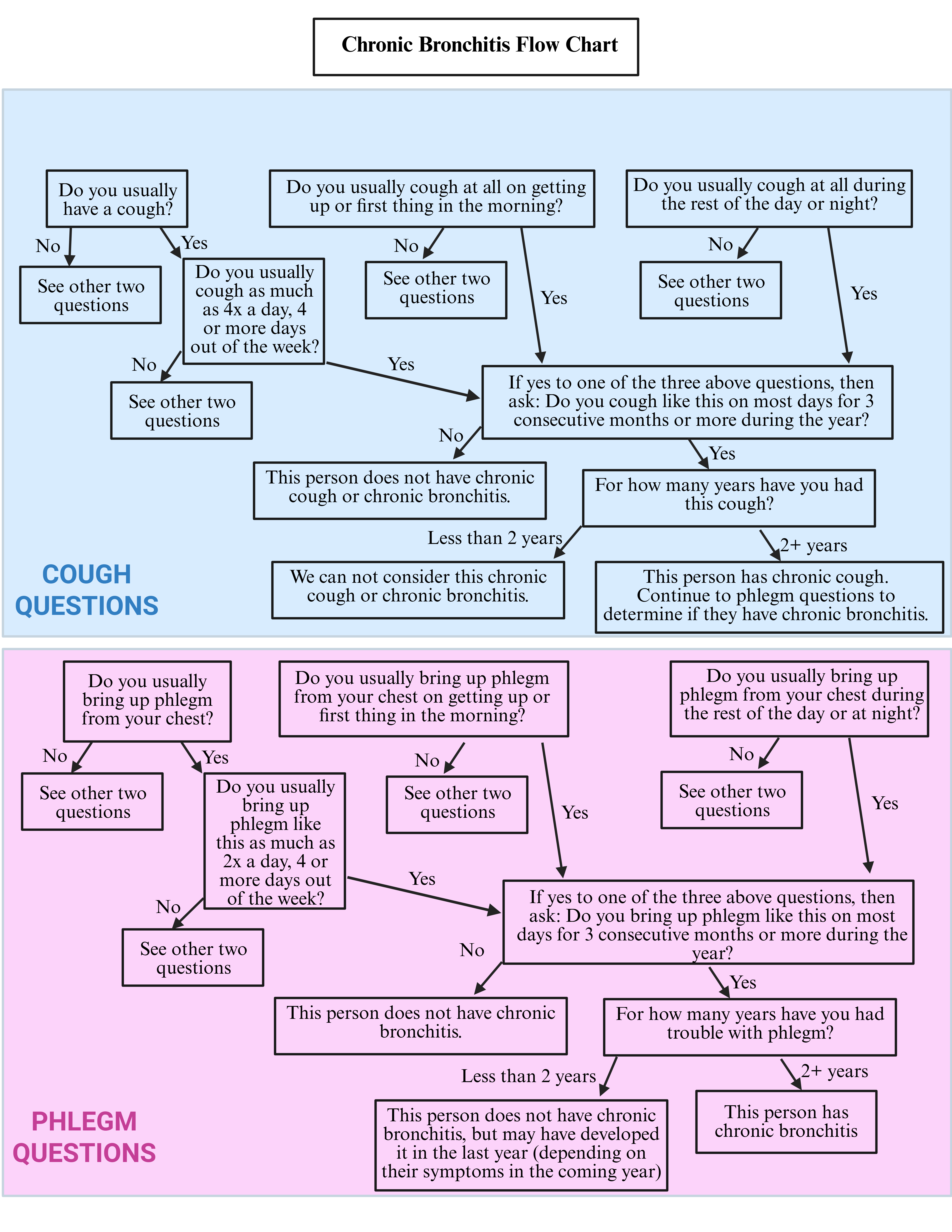

### Chronic Rhinosinusitis Flow Chart

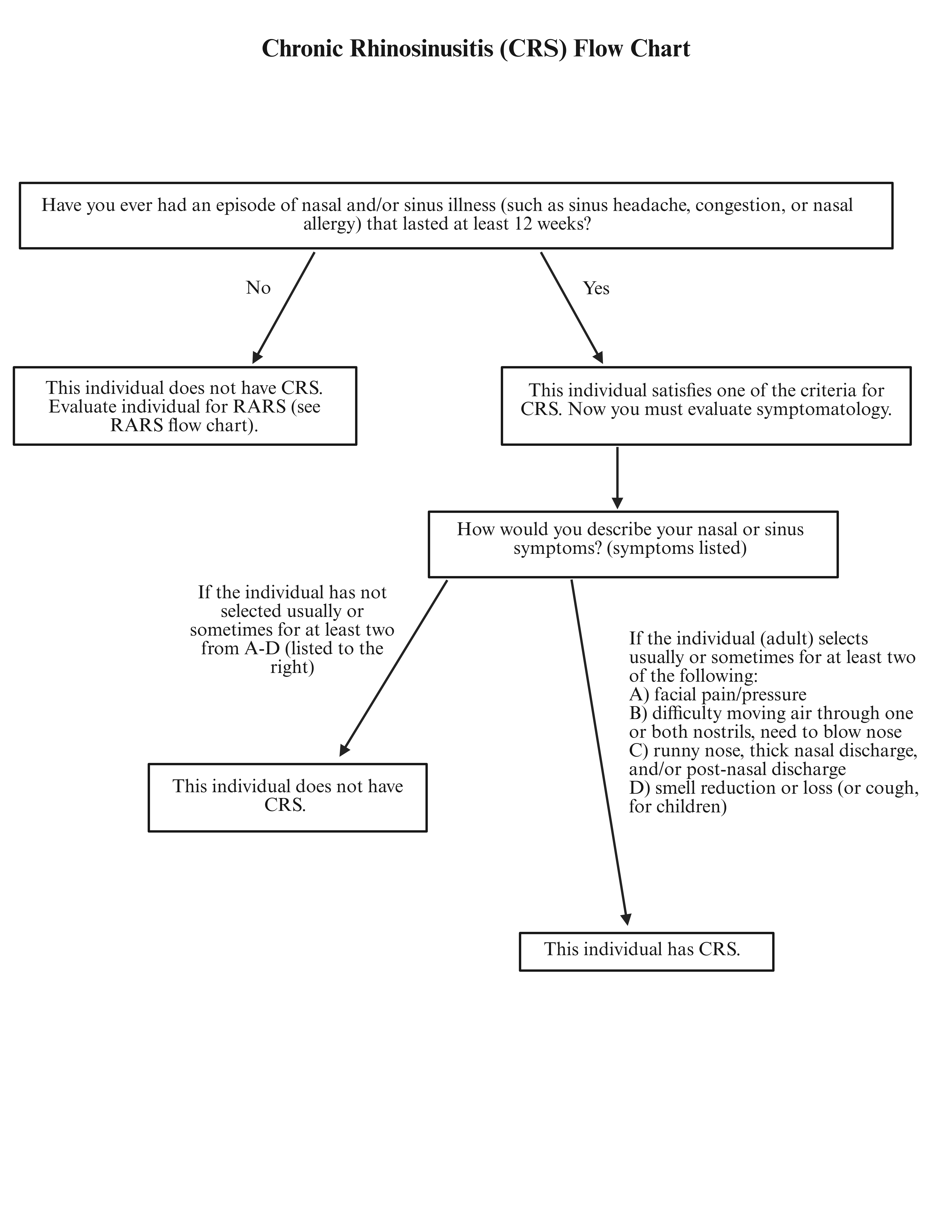

### Recurrent Acute Rhinosinusitis Flow Chart

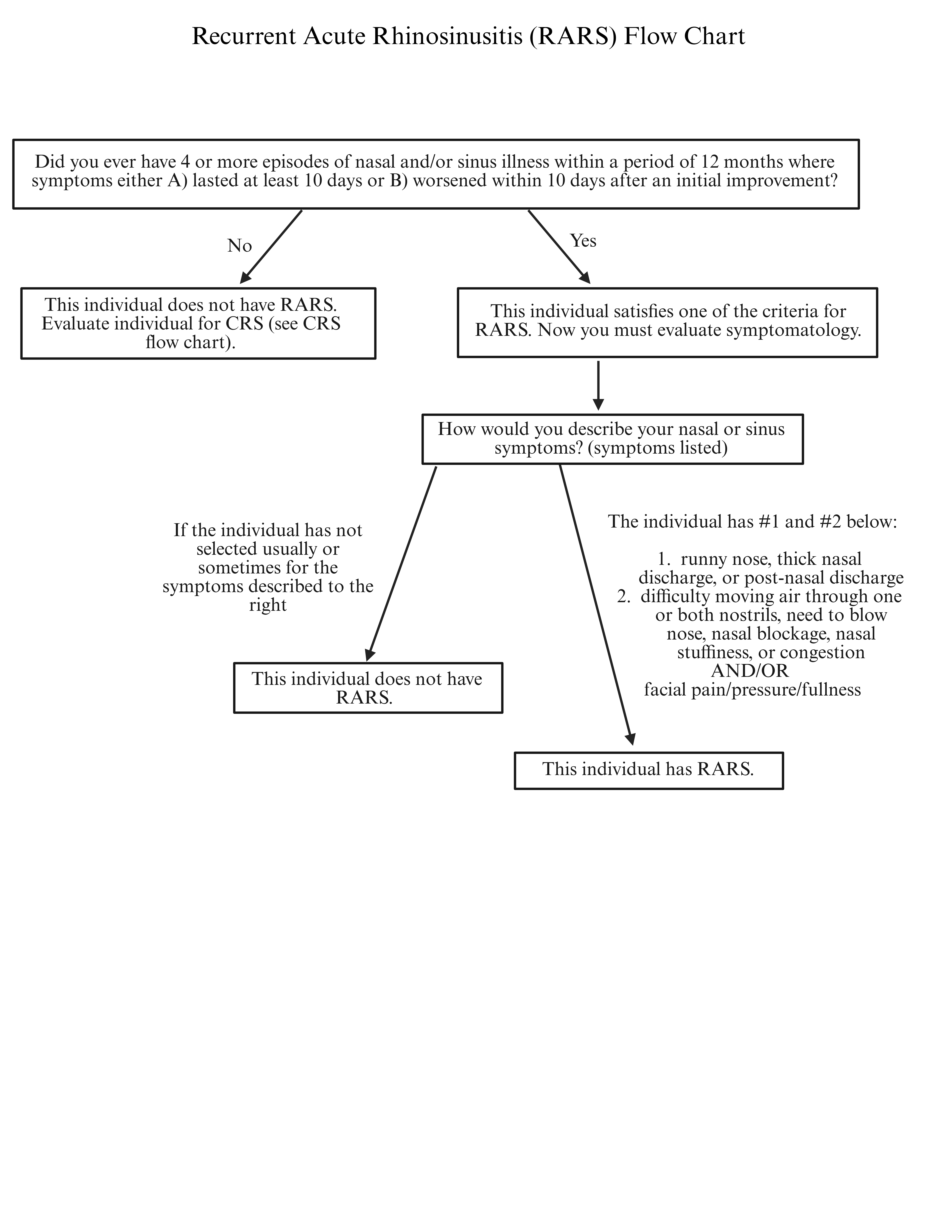

### Supplemental Figure 1

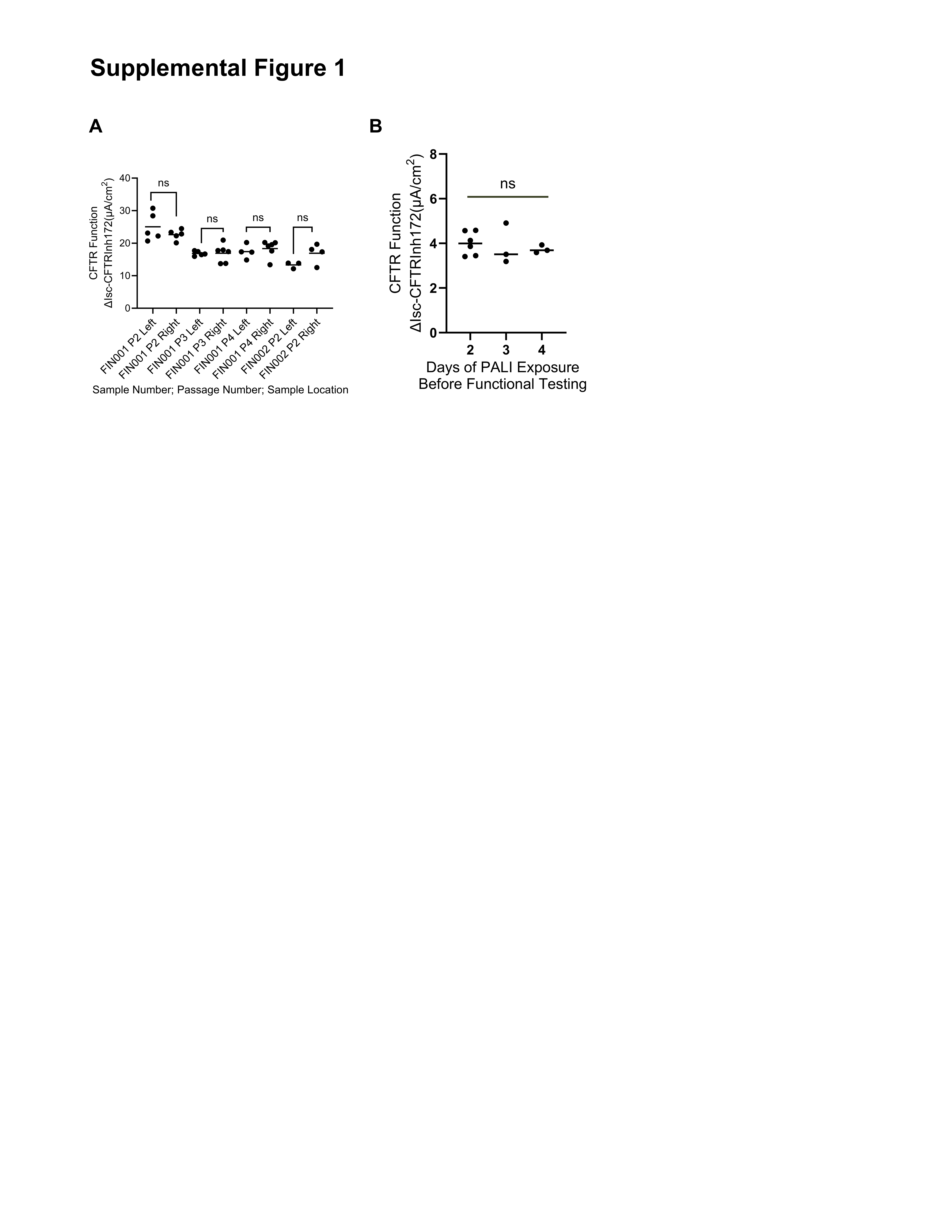

### Supplemental Figure 2

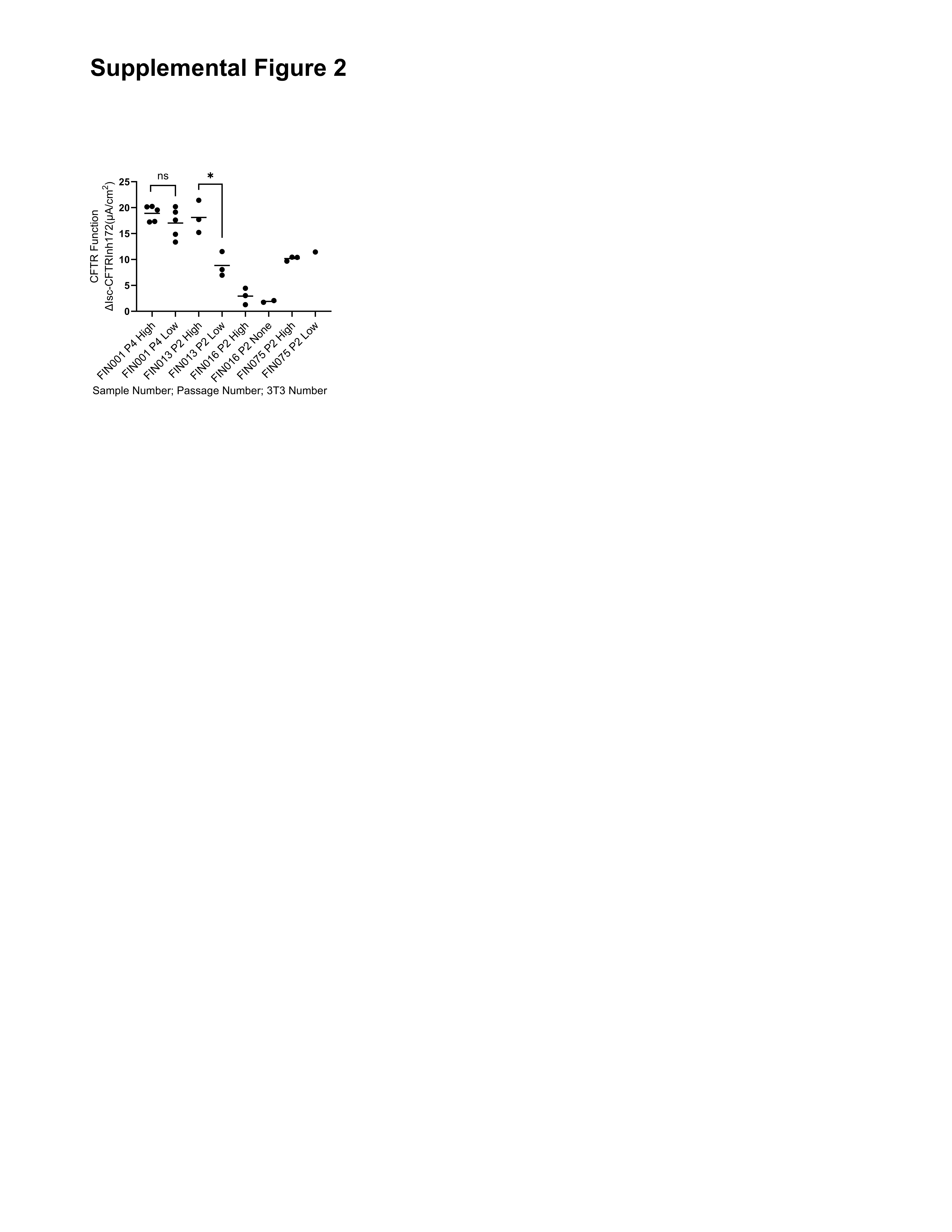

### Supplemental Figure 3

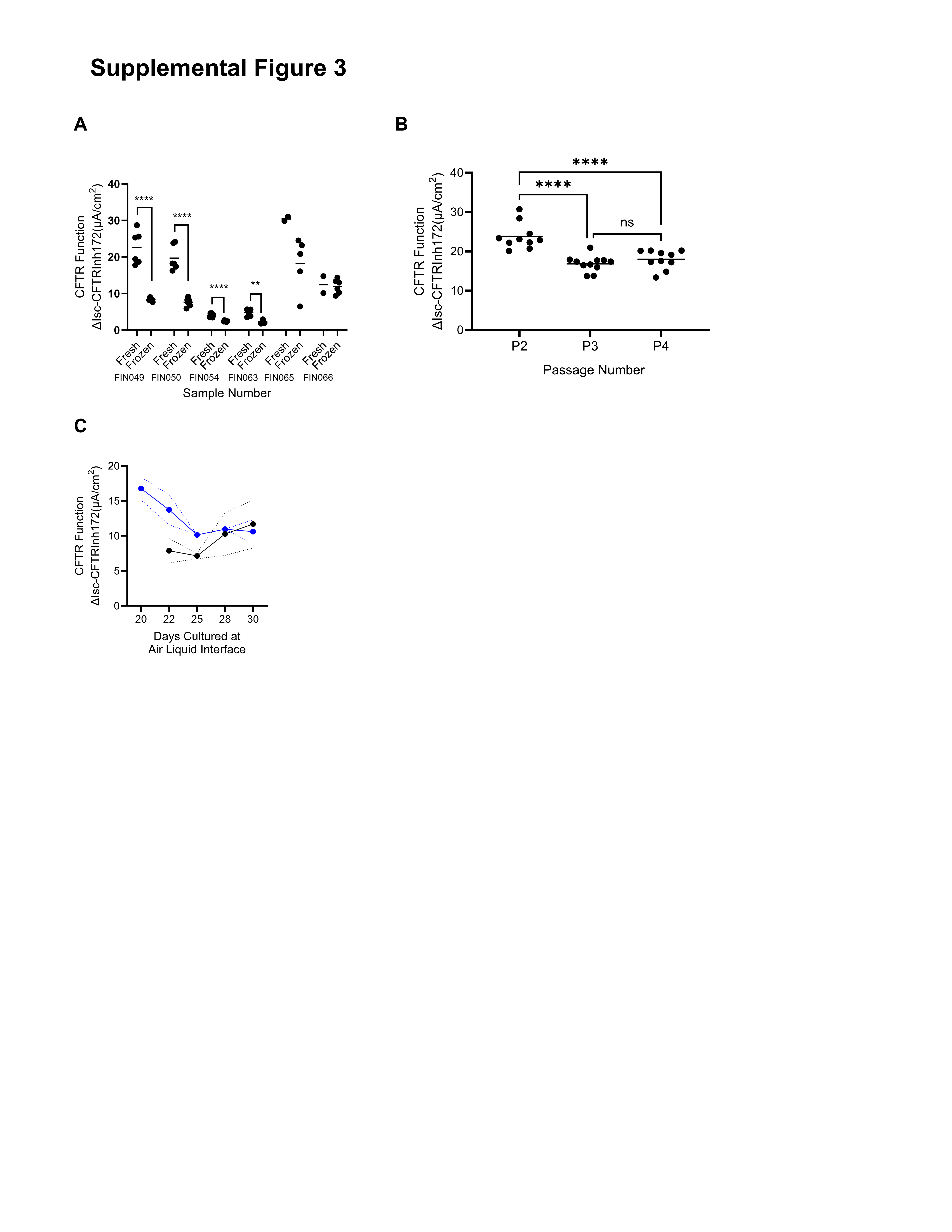

### Supplemental Figure 4

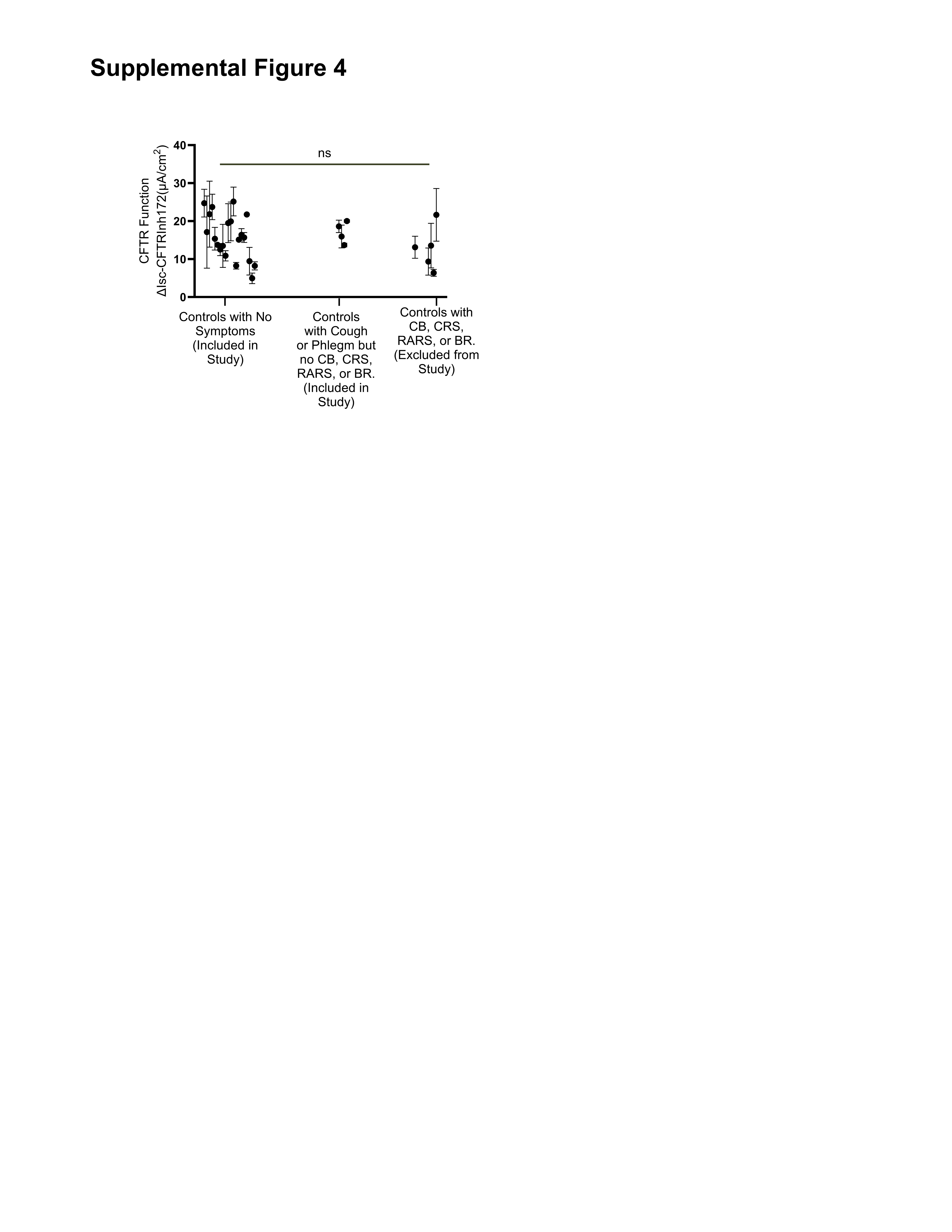

### Supplemental Figure 5

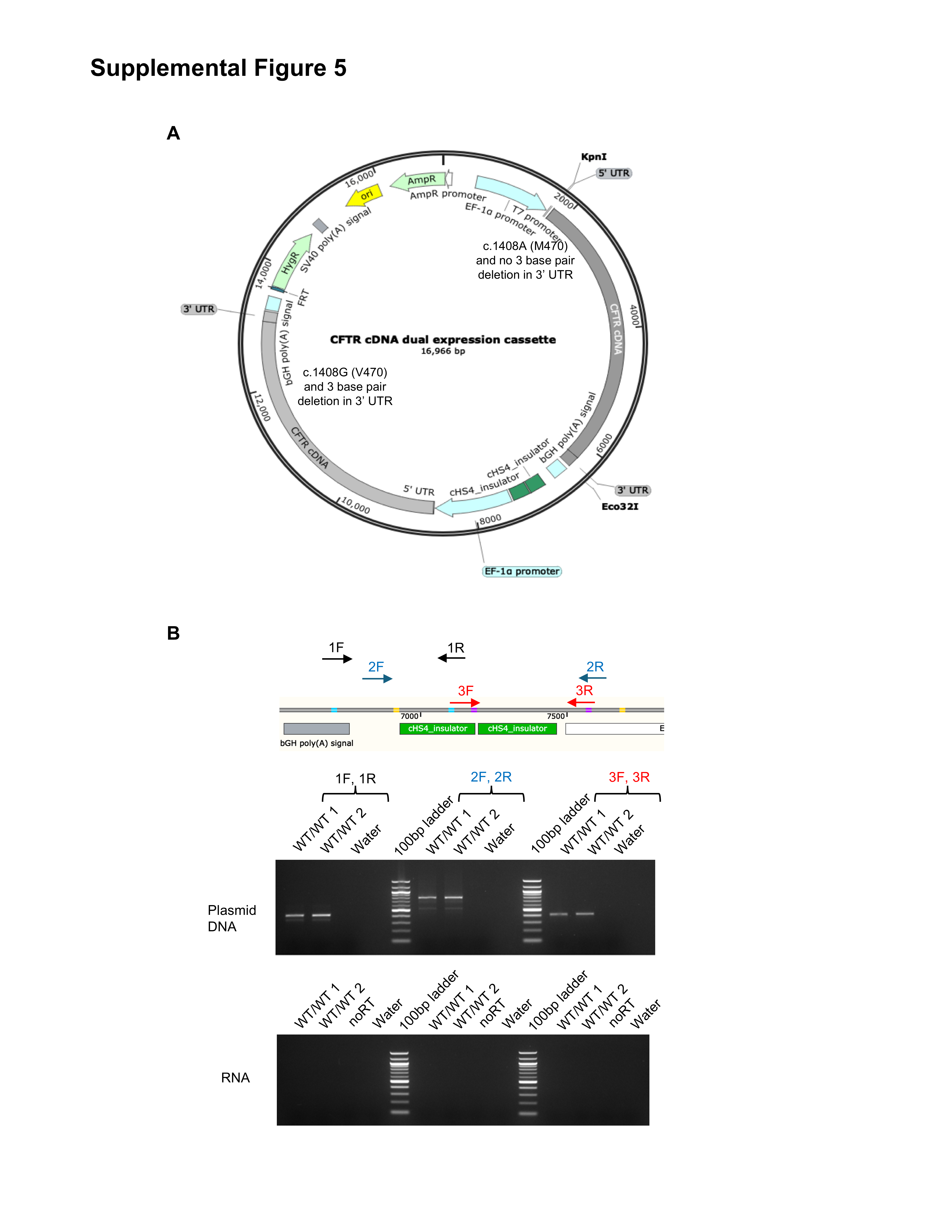

### Supplemental Figure 6

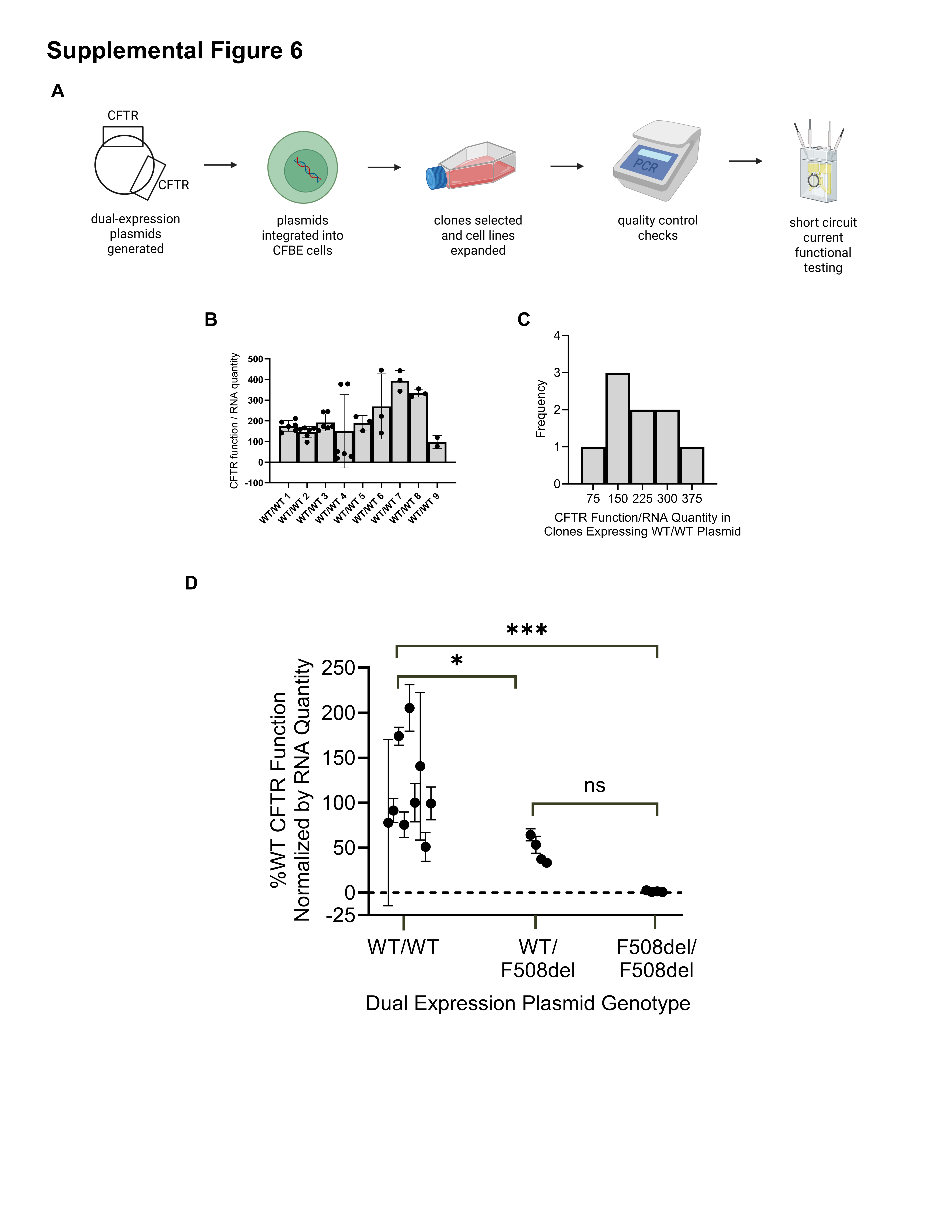

### Supplemental Figure 7

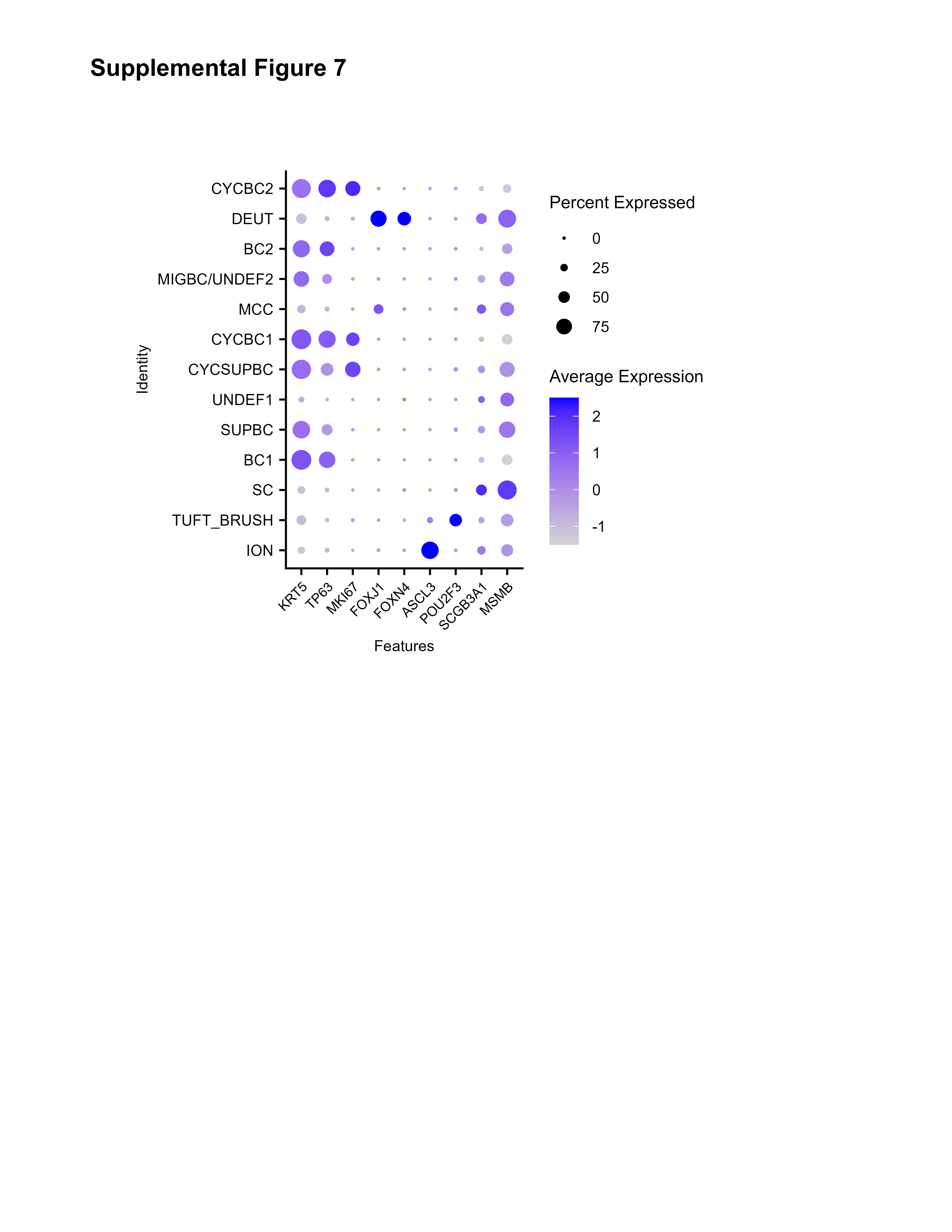
