## Supplementary material for "CFTR function in nasal airway cells from symptomatic and asymptomatic CF heterozygotes": Health History Survey

### Demographic and Health Questionnaire

Please complete the survey below.

Thank you!

#### About Me

Date completing this form (today's date) __________________________________

DEMOGRAPHICS First Name __________________________________

DEMOGRAPHICS Last Name __________________________________

DEMOGRAPHICS Email Address __________________________________

DEMOGRAPHICS Year of Birth __________________________________

DEMOGRAPHICS Sex assigned at birth __________________________________

DEMOGRAPHICS Are you left or right handed?
- ◯ Left
- ◯ Right

DEMOGRAPHICS How do you prefer we contact you if we have follow-up questions after survey completion?
- ◯ Cell phone (call)
- ◯ Cell phone (text)
- ◯ Home phone
- ◯ Email

DEMOGRAPHICS Please enter your cell phone, home phone, and/or email address: __________________________________

DEMOGRAPHICS Race
- ☐ White
- ☐ Black or African American
- ☐ Asian
- ☐ American Indian or Alaska Native
- ☐ Native Hawaiian or Other Pacific Islander
- ☐ Other
- ☐ Unknown

DEMOGRAPHICS Is the participant of Hispanic origin?
- ◯ Yes
- ◯ No
- ◯ Unknown

DEMOGRAPHICS What is your height in inches? __________________________________

DEMOGRAPHICS Current zip code (residence) __________________________________

How long have you lived at your current zip code? __________________________________________

Please provide a brief description about where you have lived in your life (state, country) and the duration of time for which you have lived in each place. Please include in this list where you were born.

__________________________________

Patient EPIC MRN (Medical Record Number) - Please leave this field blank.

__________________________________
(This field must be completed by the research team conducting the study. )

EPIC Note Reference Numbers -Please leave this field blank

__________________________________
(This field must be completed by the research team conducting the study. )

#### Health History

HEALTH Before completing the rest of the questionnaire:

Please tick in one circle to show how you describe your current health:

- ◯ Very good
- ◯ Good
- ◯ Fair
- ◯ Poor
- ◯ Very poor

HEALTH Over the last 4 weeks, I have coughed:

- ◯ Almost every day
- ◯ Several days a week
- ◯ A few days a month
- ◯ Only with lung/respiratory infections
- ◯ Not at all

HEALTH Over the last 4 weeks, I have brought up phlegm (sputum):

- ◯ Almost every day
- ◯ Several days a week
- ◯ A few days a month
- ◯ Only with lung/respiratory infections
- ◯ Not at all

CONDITIONS Has a doctor ever told you that you have chronic rhinosinusitis?

- ◯ Yes
- ◯ No
- ◯ I don't know

CONDITIONS Have you ever coughed up blood?

- ◯ Yes
- ◯ No
- ◯ I don't know

If so, please describe the frequency and volume of blood that you have coughed up.

__________________________________

CONDITIONS Have you ever been diagnosed with congenital cystic lung, otherwise known as congenital cystic adenomatoid malformation of the lung or congenital pulmonary airway malformation?

- ◯ Yes
- ◯ No
- ◯ I don't know

CONDITIONS Has a doctor ever told you that you have sinus polyps?

- ◯ Yes
- ◯ No
- ◯ I don't know

| | |
|---|---|
| CONDITIONS Has a doctor ever told you that you have respiratory failure? | ◯ Yes<br>◯ No<br>◯ I don't know |
| CONDITIONS Has a doctor ever told you that you have finger clubbing? | ◯ Yes<br>◯ No<br>◯ I don't know |

The next question will ask you to upload an image of your left or right index finger as shown in the attached image below.

The image should be taken of the side of the finger. Please make sure that you have a solid colored background. Thank you!

[Attachment: "Index_Finger_Image.jpg"]

| | |
|---|---|
| Please upload an image of your left or right index finger here: | |
| CONDITIONS Has a doctor ever told you that you have diabetes? | ◯ Yes<br>◯ No<br>◯ I don't know |
| CONDITIONS What type of diabetes have you been diagnosed with? | ◯ Type I<br>◯ Type II<br>◯ Gestational<br>◯ Prediabetes<br>◯ Maturity-onset diabetes of the young<br>◯ Secondary<br>◯ I don't know |
| CONDITIONS Has a doctor ever told you that you have pancreatitis? | ◯ Yes<br>◯ No<br>◯ I don't know |
| CONDITIONS Have you been diagnosed with acute or chronic pancreatitis? | ◯ Acute<br>◯ Chronic<br>◯ I don't know |
| CONDITIONS How many times have you had pancreatitis? | ________________________ |
| When you have had pancreatitis, how long did it last?<br><br>If you've had pancreatitis more than once, please indicate a length of time for each occurrence.<br><br>If you currently have pancreatitis, please write how long you've had it and add the word "ongoing." | ______________________________ |
| CONDITIONS Has your doctor ever provided an explanation for your pancreatitis? If so, please explain. Otherwise, please leave this question blank. | ______________________________ |
| CONDITIONS Has a doctor ever told you that you have pancreatic insufficiency? | ◯ Yes<br>◯ No<br>◯ I don't know |

| | |
|---|---|
| CONDITIONS Do you take supplemental pancreatic enzymes? For example, Creon®, Zenpep®, Pancreaze®, Ultresa®, Pertzye®, or Viokace® | ◯ Yes<br>◯ No<br>◯ I don't know |
| CONDITIONS Do you frequently experience constipation? | ◯ Yes<br>◯ No<br>◯ I don't know |
| CONDITIONS If yes, will you please define your constipation frequency. | __________________________________ |
| CONDITIONS Has a doctor ever told you that you have gastrointestinal reflux disease (GERD)? | ◯ Yes<br>◯ No<br>◯ I don't know |
| CONDITIONS Has a doctor ever told you that you have cancer? | ◯ Yes<br>◯ No<br>◯ I don't know |
| If yes, what type(s): | ____________________________ |
| CONDITIONS Has a doctor ever told you that you have a pseudomonas infection? | ◯ Yes<br>◯ No<br>◯ I don't know |
| CONDITIONS Has a doctor ever told you that you have a nontuberculous mycobacterial infection? | ◯ Yes<br>◯ No<br>◯ I don't know |
| CONDITIONS Has a doctor ever told you that you have an aspergillosis infection? | ◯ Yes<br>◯ No<br>◯ I don't know |
| CONDITIONS Has a doctor ever told you that you have a Staphylococcus aureus infection? | ◯ Yes<br>◯ No<br>◯ I don't know |
| CONDITIONS Has a doctor ever told you that you have kidney stones? | ◯ Yes<br>◯ No<br>◯ I don't know |
| CONDITIONS Has a doctor ever told you that you have hypertension (high blood pressure)? | ◯ Yes<br>◯ No<br>◯ I don't know |
| CONDITIONS Have you ever gone to a fertility doctor for reproductive needs? | ◯ Yes<br>◯ No |
| CONDITIONS Have you personally faced any difficulties in trying to conceive a child? | ◯ Yes<br>◯ No<br>◯ I have never tried to conceive<br>◯ I prefer not to answer<br>◯ I don't know |

| | |
|---|---|
| CONDITIONS Has a doctor ever told you that you have bronchiectasis? | ◯ Yes<br>◯ No<br>◯ I don't know |
| CONDITIONS Has a doctor ever told you that you have cystic fibrosis? | ◯ Yes<br>◯ No<br>◯ I don't know |
| CONDITIONS Has a doctor ever told you that you have CFTR-related disorder? | ◯ Yes<br>◯ No<br>◯ I don't know |
| CONDITIONS Has a doctor ever told you that you have CFTR-related metabolic syndrome? | ◯ Yes<br>◯ No<br>◯ I don't know |
| CONDITIONS If you are male, has a doctor ever told you that you have congenital bilateral absence of the vas deferens (CBAVD)?<br><br>If you are not male, you may skip this question. | ◯ Yes<br>◯ No<br>◯ I don't know |
| MEDICATIONS In the past year, have you been treated with antibiotics for a chest illness? | ◯ Yes<br>◯ No<br>◯ I don't know<br>(PX090901) |
| How many times? | __________________________<br>(PX090901) |
| MEDICATIONS In the past year, have you been treated with steroid pills or injections, such as prednisone or solumedrol, for a chest illness? | ◯ Yes<br>◯ No<br>◯ I don't know<br>(PX090901) |
| How many times? | __________________________<br>(PX090901) |
| MEDICATIONS Please list any medications, antibiotics, bronchodilators, corticosteroids, and CFTR modulators that you are currently using (both prescribed and over the counter). With each medication, please also indicate A) the route of administration (e.g. swallowed pill vs inhaler) and B) if the medication is being taken for pulmonary trouble. | __________________________ |
| TREATMENT Do you currently utilize any primary or secondary airway clearance techniques? (For example, positive expiratory pressure, postural drainage with clapping, forced expiratory technique, oscillating PEP, high frequency chest wall isolation.) If so, please list. Otherwise, please skip this question. | __________________________ |

| | |
|---|---|
| TREATMENT Have you used oxygen therapy in the last year? | ◯ Yes, Continuously<br>◯ Yes, Nocturnal and/or with exertion<br>◯ Yes, During exacerbation<br>◯ Yes, As needed<br>◯ No |
| CONDITIONS Have you ever contracted sars-cov-2 (COVID19)? | ◯ Yes<br>◯ No |
| CONDITIONS How many times have you contracted sars-cov-2 (COVID19)? | __________________________________ |
| CONDITIONS If you frequently experience nasal and sinus symptoms, do you think that your symptoms have worsened since contracting sars-cov-2 (COVID-19) e.g. long covid? | ☐ my symptoms have worsened since contracting COVID-19<br>☐ my symptoms are still the same as they were before contracting COVID-19<br>☐ I did not have sinus and nasal symptoms before contracting COVID-19, but now I experience symptoms frequently<br>☐ I don't experience frequent nasal and sinus symptoms<br>☐ I don't know |

### Chronic Rhinosinusitis Questionnaire

NASAL/SINUS Have you ever had an episode of nasal and/or sinus illness (such as sinus headache, congestion, or nasal allergy) that lasted for more than 12 weeks?

◯ Yes
◯ No

NASAL/SINUS Did you ever have 4 or more episodes of nasal and/or sinus illness within a period of 12 months where symptoms either A) lasted at least 10 days or B) worsened within 10 days after an initial improvement?

◯ Yes
◯ No

NASAL/SINUS Have you ever used medication or other methods to prevent or treat nasal and/or sinus trouble other than treatment for cold and flu?

◯ Yes
◯ No

NASAL/SINUS If you have (or had) a nasal and/or sinus problem, how old were you when the problem was first noted? (Please specify your answer by entering a number followed by a unit. For example, if you first noted a sinus problem at 21 years old, please type 21 years.)

__________________________________

NASAL/SINUS If you have had a nasal and/or sinus problem, has the problem resolved for you?

◯ Yes
◯ No

NASAL/SINUS At what age did your nasal and/or sinus problem resolve? (Please specify your answer by entering a number followed by a unit. For example, if you first noted a sinus problem at 21 years old, please type 21 years.)

__________________________________

#### Advanced Questionnaire for Chronic Sinusitis Study

NASAL/SINUS Have you ever been told by a doctor that you have nasal allergies?

◯ Yes
◯ No

NASAL/SINUS If you have had separate episodes of nasal or sinus problems each year, do these episodes seem to be in the same season(s) each year?

◯ Yes
◯ No

If yes, which one(s):

☐ Spring
☐ Summer
☐ Fall
☐ Winter

NASAL/SINUS If you don't think your nasal or sinus problems have any seasonal pattern, can you think of any triggers for the episodes?

Always no reason ______

Cold or infection ______

Change in weather ______

Exposure to smoke or air pollution ______

Exposure to pets ______

Some foods ______

Others (Please list) ______

| | |
|---|---|
| CONDITIONS Have you ever been told by a doctor that you have asthma? | ◯ Yes<br>◯ No |
| CONDITIONS Have you ever been told by a doctor that you have any chronic lung disease? | ◯ Yes<br>◯ No |
| If yes, please specify | ______________________ |

NASAL/SINUS How would you describe your nasal or sinus symptoms?

Facial pain / pressure / fullness

______

Reduction or loss of sense of smell

______

Difficulty moving air through one or both nostrils

______

Nasal blockage

______

Nasal stuffiness

______

Congestion

______

Ear fullness / pain

______

Fatigue

______

Dizziness

______

Cough

______

Sneezing

______

Runny nose

______

Itching of nose, palate, or throat

______

Need to blow nose

______

Thick nasal discharge

______

Post-nasal discharge

______

Reduced productivity/concentration

______

Headache

______

Partial or full loss of the sense of smell

| | |
|---|---|
| NASAL/SINUS Which are the three most disturbing symptoms above (in order)? (Please number your responses as 1., 2., 3. and make response #1 be your most disturbing symptom, followed by response #2 as your second most disturbing symptom and so on.) | __________________________________ |
| Is your nasal discharge cloudy or colored? | ◯ Yes, my nasal discharge is cloudy or colored<br>◯ No, my nasal discharge is NOT cloudy or colored<br>◯ I don't know<br>◯ I don't have nasal discharge |
| NASAL/SINUS What do you use to treat your nasal or sinus problem (all applied)? | __________________________________ |
| CONDITIONS Have you ever been told by a doctor that you have nasal polyps? | ◯ Yes<br>◯ No |
| CONDITIONS Have you ever had sinus surgery? | ◯ Yes<br>◯ No |
| CONDITIONS If yes, how many sinus surgeries have you had? | ____________________________ |
| CONDITIONS Do you frequently have problems with digestion? | ◯ Yes<br>◯ No |
| CONDITIONS Do you frequently have greasy stools or stools that float? | ◯ Yes<br>◯ No |

FAMILY_HISTORY Could you describe your grandpa and your grandma's origins here? (For example, if your father's parents have German/Japanese ancestry and your mother's parents have Italian/African-American ancestry, please write "father's parents: German/Japanese, mother's parents: Italian/African-American")

Father's parents: ______
Mother's parents: ______

Please list your children with their name, age, and sex and tell us who has cystic fibrosis (CF) and who has sinus problems similar to yours.

Name Age Sex CF/Sinus problem/None
______ ______ ______ ______
______ ______ ______ ______
______ ______ ______ ______
______ ______ ______ ______
______ ______ ______ ______
______ ______ ______ ______
______ ______ ______ ______
______ ______ ______ ______
______ ______ ______ ______

### ATS-DLD Phenx History Of Respiratory Symptoms

SYMPTOMS

These questions pertain mainly to your chest. Please answer yes or no, if possible. If a question does not appear to be applicable to you, check the "Does Not Apply" space or leave the question blank. If you are in doubt about whether your answer is yes or no, record no.

| | |
|---|---|
| COUGH Do you usually have a cough? (Count a cough with first smoke or on first going out-of-doors. Exclude clearing of throat.) | ◯ Yes<br>◯ No<br>(PX090901) |
| COUGH Do you usually cough as much as 4 to 6 times a day, 4 or more days out of the week? | ◯ Yes<br>◯ No<br>(PX090901) |
| COUGH Do you usually cough at all on getting up, or first thing in the morning? | ◯ Yes<br>◯ No<br>(PX090901) |
| COUGH Do you usually cough at all during the rest of the day or at night? | ◯ Yes<br>◯ No<br>(PX090901) |
| COUGH Do you usually cough like this on most days for 3 consecutive months or more during the year? | ◯ Yes<br>◯ No<br>◯ Does not apply<br>(PX090901) |
| COUGH For how many years have you had this cough? | __________________________________<br>(PX090901) |
| PHLEGM Do you usually bring up phlegm from your chest? (Count phlegm with the first smoke or on first going out-of-doors. Exclude phlegm from the nose. Count swallowed phlegm.) | ◯ Yes<br>◯ No<br>(PX090901) |
| PHLEGM Do you usually bring up phlegm like this as much as twice a day, 4 or more days out of the week? | ◯ Yes<br>◯ No<br>(PX090901) |
| PHLEGM Do you usually bring up phlegm at all on getting up or first thing in the morning? | ◯ Yes<br>◯ No<br>(PX090901) |
| PHLEGM Do you usually bring up phlegm at all during the rest of the day or at night? | ◯ Yes<br>◯ No<br>(PX090901) |
| PHLEGM Do you bring up phlegm like this on most days for 3 consecutive months or more during the year? | ◯ Yes<br>◯ No<br>◯ Does not apply<br>(PX090901) |
| PHLEGM For how many years have you had trouble with phlegm? | __________________________________<br>(PX090901) |

| | |
|---|---|
| EPISODES OF COUGH AND PHLEGM Have you had periods or episodes of (increased*) cough and phlegm lasting for 3 weeks or more each year? (*For individuals who usually have cough and/or phlegm) | ◯ Yes<br>◯ No<br>(PX090901) |
| EPISODES OF COUGH AND PHLEGM For how long have you had at least 1 such episode per year? | ________________________<br>(PX090901) |
| EPISODES OF COUGH AND PHLEGM About how many such episodes have you had in the past 12 months? | ________________________<br>(PX090901) |
| CHEST COLDS AND CHEST ILLNESSES If you get a cold, does it usually go to your chest? (Usually means more than 1/2 the time.) | ◯ Yes<br>◯ No<br>◯ Don't get colds<br>(PX090901) |
| CHEST COLDS AND CHEST ILLNESSES During the past 3 years, have you had any chest illnesses that have kept you off work, indoors at home, or in bed? | ◯ Yes<br>◯ No<br>(PX090901) |
| CHEST COLDS AND CHEST ILLNESSES Did you produce phlegm with any of these chest illnesses? | ◯ Yes<br>◯ No<br>◯ Does not apply<br>(PX090901) |
| CHEST COLDS AND CHEST ILLNESSES In the last 3 years, how many such illnesses, with (increased) phlegm, did you have which lasted a week or more? Number of illnesses: | ________________________<br>(PX090901) |
| PAST ILLNESSES - Lung Trouble Did you have any lung trouble before the age of 16? | ◯ Yes<br>◯ No<br>(PX090901) |
| PAST ILLNESSES - Bronchitis Have you ever had attacks of bronchitis? | ◯ Yes<br>◯ No<br>(PX090901) |
| PAST ILLNESSES - Bronchitis Was it confirmed by a doctor? | ◯ Yes<br>◯ No<br>◯ Does not apply<br>(PX090901) |
| PAST ILLNESSES - Bronchitis At what age was your first attack? | ________________________<br>(PX090901) |
| PAST ILLNESSES - Pneumonia Have you ever had Pneumonia (include bronchopneumonia)? | ◯ Yes<br>◯ No<br>(PX090901) |
| PAST ILLNESSES - Pneumonia Was it confirmed by a doctor? | ◯ Yes<br>◯ No<br>◯ Does not apply<br>(PX090901) |

| | |
|---|---|
| PAST ILLNESSES - Pneumonia At what age did you first have it? | __________________<br>(PX090901) |
| PAST ILLNESSES - Chronic Bronchitis Have you ever had chronic bronchitis? | ◯ Yes<br>◯ No<br>(PX090901) |
| PAST ILLNESSES - Chronic Bronchitis Do you still have it? | ◯ Yes<br>◯ No<br>◯ Does not apply<br>(PX090901) |
| PAST ILLNESSES - Chronic Bronchitis Was it confirmed by a doctor? | ◯ Yes<br>◯ No<br>◯ Does not apply<br>(PX090901) |
| PAST ILLNESSES - Chronic Bronchitis At what age did it start? | __________________<br>(PX090901) |
| PAST ILLNESSES - Emphysema Have you ever had emphysema? | ◯ Yes<br>◯ No<br>(PX090901) |
| PAST ILLNESSES - Emphysema Do you still have it? | ◯ Yes<br>◯ No<br>◯ Does not apply<br>(PX090901) |
| PAST ILLNESSES - Emphysema Was it confirmed by a doctor? | ◯ Yes<br>◯ No<br>◯ Does not apply<br>(PX090901) |
| PAST ILLNESSES - Emphysema At what age did it start? | __________________<br>(PX090901) |
| PAST ILLNESSES - Asthma Have you ever had asthma? | ◯ Yes<br>◯ No<br>(PX090901) |
| PAST ILLNESSES - Asthma Do you still have it? | ◯ Yes<br>◯ No<br>◯ Does not apply<br>(PX090901) |
| PAST ILLNESSES - Asthma Was it confirmed by a doctor? | ◯ Yes<br>◯ No<br>◯ Does not apply<br>(PX090901) |
| PAST ILLNESSES - Asthma At what age did it start? | __________________<br>(PX090901) |

| | |
|---|---|
| PAST ILLNESSES - Asthma If you no longer have it, at what age did it stop? | __________________________________ (PX090901) |
| PAST ILLNESSES - Other Chest Illnesses Have you ever had Any other chest illnesses? | ◯ Yes<br>◯ No<br>(PX090901) |
| PAST ILLNESSES - Other Chest Illnesses Please specify chest illnesses you had. | __________________________________ (PX090901) |
| OCCUPATIONAL HISTORY - Dusty Job Have you ever worked for a year or more in any dusty job? | ◯ Yes<br>◯ No<br>◯ Does not apply<br>(PX090901) |
| OCCUPATIONAL HISTORY - Dusty Job Specify job/industry: | __________________________________ (PX090901) |
| OCCUPATIONAL HISTORY - Dusty Job Total years worked? | __________________________________ (PX090901) |
| OCCUPATIONAL HISTORY - Dusty Job Was dust exposure mild, moderate, or severe? | ◯ Mild<br>◯ Moderate<br>◯ Severe<br>(PX090901) |
| OCCUPATIONAL HISTORY - Gas or Chemical Fumes Have you ever been exposed to gas or chemical fumes in your work? | ◯ Yes<br>◯ No<br>◯ Does not apply<br>(PX090901) |
| OCCUPATIONAL HISTORY - Gas or Chemical Fumes Specify job/industry: | __________________________________ (PX090901) |
| OCCUPATIONAL HISTORY - Gas or Chemical Fumes Total years worked? | __________________________________ (PX090901) |
| OCCUPATIONAL HISTORY - Gas or Chemical Fumes Was gas or chemical fumes exposure mild, moderate, or severe? | ◯ Mild<br>◯ Moderate<br>◯ Severe<br>(PX090901) |
| TOBACCO SMOKING Have you ever smoked cigarettes? (NO means less than 20 packs of cigarettes or 12 oz. of tobacco in a lifetime or less than 1 cigarette a day for 1 year.) | ◯ Yes<br>◯ No<br>(PX090901) |
| TOBACCO SMOKING Do you now smoke cigarettes (as of 1 month ago)? | ◯ Yes<br>◯ No<br>◯ Does not apply<br>(PX090901) |

| | |
|---|---|
| TOBACCO SMOKING How old were you when you first started regular cigarette smoking? | ____________________ (PX090901) |
| TOBACCO SMOKING If you have stopped smoking cigarettes completely, how old were you when you stopped? Age stopped: | ____________________ (PX090901) |
| TOBACCO SMOKING How many cigarettes do you smoke per day now? | ____________________ (PX090901) |
| TOBACCO SMOKING On the average of the entire time you smoked, how many cigarettes did you smoke per day? | ____________________ (PX090901) |
| TOBACCO SMOKING Do or did you inhale the cigarette smoke? | ◯ Does not apply<br>◯ Not at all<br>◯ Slightly<br>◯ Moderately<br>◯ Deeply<br>(PX090901) |
| TOBACCO SMOKING Have you ever been exposed to secondhand cigarette smoke? | ◯ Yes<br>◯ No |
| TOBACCO SMOKING If yes to the previous question, please describe the frequency and circumstances surrounding your cigarette smoke exposure. | ____________________ |
| TOBACCO SMOKING Have you ever smoked a pipe regularly? (YES means more than 12 oz. tobacco in a lifetime.) | ◯ Yes<br>◯ No<br>(PX090901) |
| TOBACCO SMOKING How old were you when you started to smoke a pipe regularly? Age: | ____________________ (PX090901) |
| TOBACCO SMOKING If you have stopped smoking a pipe completely, how old were you when you stopped? Age stopped: | ____________________ (PX090901) |
| TOBACCO SMOKING On the average over the entire time you smoked a pipe, how much pipe tobacco did you smoke per week? Oz. per week (a standard pouch of tobacco contains 1 1/2 oz.): | ____________________ (PX090901) |
| TOBACCO SMOKING How much pipe tobacco are you smoking now? | ____________________ (PX090901) |
| TOBACCO SMOKING Do or did you inhale the pipe smoke? | ◯ Never smoked<br>◯ Not at all<br>◯ Slightly<br>◯ Moderately<br>◯ Deeply<br>(PX090901) |

| | |
|---|---|
| TOBACCO SMOKING Have you ever smoked cigars regularly? (YES means more than 1 cigar a week for a year.) | ◯ Yes<br>◯ No<br>(PX090901) |
| TOBACCO SMOKING How old were you when you started smoking cigars regularly? Age: | ______________________<br>(PX090901) |
| TOBACCO SMOKING If you have stopped smoking cigars completely, how old were you when you stopped? Age stopped: | ______________________<br>(PX090901) |
| TOBACCO SMOKING On the average over the entire time you smoked cigars, how many cigars did you smoke per week? | ______________________<br>(PX090901) |
| TOBACCO SMOKING How many cigars are you smoking per week now? | ______________________<br>(PX090901) |
| TOBACCO SMOKING Do or did you inhale the cigar smoke? | ◯ Never smoked<br>◯ Not at all<br>◯ Slightly<br>◯ Moderately<br>◯ Deeply<br>(PX090901) |
| MARIJUANA Have you ever used marijuana in any form? | ◯ Yes<br>◯ No<br>◯ I don't know |
| MARIJUANA Have you ever smoked marijuana? | ◯ Yes<br>◯ No<br>◯ I don't know |
| MARIJUANA Which category best describes the total number of times you've smoked marijuana over your lifetime? | ◯ 0-50<br>◯ 51-500<br>◯ 501-1000<br>◯ More than 1000<br>◯ I don't know |
| MARIJUANA Over the entire period you smoked marijuana, how many years did you smoke marijuana on a daily or near-daily basis? | ______________________ |
| MARIJUANA Have you smoked marijuana in the last 30 days? | ◯ Yes<br>◯ No<br>◯ I don't know |
| MARIJUANA In the last 30 days, have you smoked a: | ☐ Joint<br>☐ Pipe<br>☐ Bong<br>☐ Blunt<br>☐ Spliff<br>☐ I don't know<br>(Definitions: A blunt is a hollowed-out cigar filled with marijuana. A spliff is a marijuana cigarette prepared with both marijuana and tobacco.) |

| | |
|---|---|
| MARIJUANA In the last 30 days, have you combined tobacco and marijuana to smoke at a single time? | ◯ Yes<br>◯ No<br>◯ I don't know |
| MARIJUANA In the last 30 days, how many days per week did you smoke a blunt? | ◯ Every day<br>◯ 6 days/week<br>◯ 5 days/week<br>◯ 4 days/week<br>◯ 3 days/week<br>◯ 2 days/week<br>◯ 1 day/week<br>◯ ______ Day(s)/month<br>◯ I don't know |
| MARIJUANA In the last 30 days, how many days per week did you smoke a spliff? | ◯ Every day<br>◯ 6 days/week<br>◯ 5 days/week<br>◯ 4 days/week<br>◯ 3 days/week<br>◯ 2 days/week<br>◯ 1 day/week<br>◯ ______ Day(s)/month<br>◯ I don't know |
| MARIJUANA In the last 30 days, how many days per week did you smoke a joint? | ◯ Every day<br>◯ 6 days/week<br>◯ 5 days/week<br>◯ 4 days/week<br>◯ 3 days/week<br>◯ 2 days/week<br>◯ 1 day/week<br>◯ ______ Day(s)/month<br>◯ I don't know |
| MARIJUANA On the days you smoked in the last 30 days, how many joints did you smoke a day? | __________________________________ |
| MARIJUANA In the last 30 days, how many days per week did you smoke a pipe? | ◯ Every day<br>◯ 6 days/week<br>◯ 5 days/week<br>◯ 4 days/week<br>◯ 3 days/week<br>◯ 2 days/week<br>◯ 1 day/week<br>◯ ______ Day(s)/month<br>◯ I don't know |
| MARIJUANA On the days you smoked in the last 30 days, how many pipes did you smoke a day? | __________________________________ |
| MARIJUANA In the last 30 days, how many days per week did you smoke a bong? | ◯ Every day<br>◯ 6 days/week<br>◯ 5 days/week<br>◯ 4 days/week<br>◯ 3 days/week<br>◯ 2 days/week<br>◯ 1 day/week<br>◯ ______ Day(s)/Month<br>◯ I don't know |

| | |
|---|---|
| MARIJUANA On the days you smoked in the last 30 days, how many bongs did you smoke a day? | ________________________________ |
| MARIJUANA Which of the following categories best captures the amount of marijuana you smoked over the past 30 days? | ◯ ______ Hit(s)/month<br>◯ ______ Gram(s)/month<br>◯ An eighth of an ounce, which is the same as 3.5 grams<br>◯ A quarter of an ounce, which is the same as 7 grams<br>◯ A half of an ounce, which is the same as 14 grams<br>◯ Three quarters of an ounce, which is the same as 21 grams<br>◯ An ounce, which is the same as 28 grams<br>◯ ______ Ounce(s)/month<br>◯ Other measure: ______<br>◯ I don't know |
| MARIJUANA Have you ever vaporized marijuana? | ◯ Yes<br>◯ No<br>◯ I don't know<br>(Vaping is the act of inhaling vapor produced by a vaporizer or electronic marijuana cigarette.) |
| MARIJUANA Which category best describes the total number of times you've vaporized marijuana over your lifetime? | ◯ 0-50<br>◯ 51-500<br>◯ 501-1000<br>◯ More than 1000<br>◯ I don't know |
| MARIJUANA Over the period you vaporized marijuana, how many years did you vaporize marijuana on a daily or near-daily basis? | ________________________________ |
| MARIJUANA Have you vaporized marijuana in the last 30 days? | ◯ Yes<br>◯ No<br>◯ I don't know |
| MARIJUANA In the last 30 days, how many days per week did you vaporize marijuana? | ◯ Every day<br>◯ 6 days/week<br>◯ 5 days/week<br>◯ 4 days/week<br>◯ 3 days/week<br>◯ 2 days/week<br>◯ 1 day/week<br>◯ ______ Day(s)/Month<br>◯ I don't know |
| MARIJUANA Have you ever dabbed marijuana? | ◯ Yes<br>◯ No<br>◯ I don't know<br>(Dabbing is a way of heating butane hash oil to extremely high temperatures to inhale the fumes.) |
| MARIJUANA Which category best describes the total number of times you've dabbed marijuana over your lifetime? | ◯ 0-50<br>◯ 51-500<br>◯ 501-1000<br>◯ More than 1000<br>◯ I don't know |

| | |
|---|---|
| MARIJUANA Have you dabbed marijuana in the last 30 days? | ◯ Yes<br>◯ No<br>◯ I don't know |
| MARIJUANA In the last 30 days, how many days per week did you dab marijuana? | ◯ Every day<br>◯ 6 days/week<br>◯ 5 days/week<br>◯ 4 days/week<br>◯ 3 days/week<br>◯ 2 days/week<br>◯ 1 day/week<br>◯ ______ Day(s)/Month<br>◯ I don't know |
| ELECTRONIC CIGARETTES Have you ever used a vaping product, even one time? | ◯ Yes<br>◯ No<br>◯ I have never heard of vaping products |
| ELECTRONIC CIGARETTES At the time when you were vaping most often, how often did you vape? | ◯ Daily<br>◯ Less than daily, but at least once a week<br>◯ Less than weekly, but at least occasionally<br>◯ I have only tried vaping a few times, but more than once<br>◯ I have only ever tried vaping once |
| ELECTRONIC CIGARETTES Have you ever vaped at least weekly, or always less than weekly? | ◯ At least weekly<br>◯ Less than weekly |
| ELECTRONIC CIGARETTES Thinking back to 24 months ago, how often were you vaping? | ◯ Daily<br>◯ Less than daily, but at least once a week<br>◯ Less than weekly, but at least once a month<br>◯ Less than monthly, but occasionally<br>◯ Not at all |
| ELECTRONIC CIGARETTES Choose the nicotine strength of the e-liquid, cartridges, or pods you currently use most / last used? | ◯ None (0mg/ml nicotine)<br>◯ Less than 10mg/ml<br>◯ 10-19 mg/ml<br>◯ 20-29 mg/ml<br>◯ 30-39 mg/ml<br>◯ 40-49 mg/ml<br>◯ 50 mg/ml or more |
| ELECTRONIC CIGARETTES When was the last time you vaped? | ◯ Less than 1 week ago<br>◯ 1-4 weeks ago<br>◯ 1-3 months ago<br>◯ 4-6 months ago<br>◯ 7-12 months ago<br>◯ 13-18 months ago<br>◯ 19-24 months ago<br>◯ More than 2 years ago |
| ELECTRONIC CIGARETTES How often, if at all, do you currently use vaping products (i.e., vape)? | ◯ Daily<br>◯ Less than daily, but at least once a week<br>◯ Less than weekly, but at least once a month<br>◯ Less than once a month, but occasionally<br>◯ Not at all |

| | |
|---|---|
| ELECTRONIC CIGARETTES In a typical week, on how many days do you vape? | ◯ 1-2 days a week<br>◯ 3-4 days a week<br>◯ 5-6 days a week<br>◯ Can't say, there's no consistent pattern |
| ELECTRONIC CIGARETTES For how long have you been vaping at least once a week? | ◯ Less than 1 month<br>◯ 1-3 months<br>◯ 4-6 months<br>◯ 7-12 months<br>◯ 13-18 months<br>◯ 19-24 months<br>◯ 2-3 years<br>◯ 3-5 years<br>◯ More than 5 years |
| ELECTRONIC CIGARETTES For how long have you been vaping daily? | ◯ Less than 1 week<br>◯ 1-4 weeks<br>◯ 1-3 months<br>◯ 4-6 months<br>◯ 7-12 months<br>◯ 13-18 months<br>◯ 19-24 months<br>◯ 2-3 years<br>◯ 3-5 years<br>◯ More than 5 years |
| ELECTRONIC CIGARETTES Which of the following e-liquid flavor categories have you used in the past 30 days? | ☐ Tobacco flavor<br>☐ Mix of tobacco and menthol<br>☐ Menthol or mint<br>☐ Fruit flavor<br>☐ desserts, sweets<br>☐ Chocolate<br>☐ Clove or other spice<br>☐ Coffee<br>☐ A non-alcoholic drink (soda, energy drinks, or other beverages)<br>☐ An alcoholic drink (wine, whiskey, cognac, margarita, or other cocktails)<br>☐ Unflavored e-liquid |
| FAMILY HISTORY Was your biological father ever told by a doctor that he has any of the following chronic lung conditions: | ☐ Chronic bronchitis<br>☐ Emphysema<br>☐ Asthma<br>☐ Lung Cancer<br>☐ Other<br>(PX090901) |
| If other, please list: | __________________________________ |
| FAMILY HISTORY Was your biological mother ever told by a doctor that she has any of the following chronic lung conditions: | ☐ Chronic bronchitis<br>☐ Emphysema<br>☐ Asthma<br>☐ Lung Cancer<br>☐ Other<br>(PX090901) |
| If other, please list: | __________________________________ |

In the past year, have you been to the emergency room or hospitalized for lung problems?

◯ Yes
◯ No
(PX090901)

How many times?

__________________________________
(PX090901)
